# Quantifying Cellular Capacity to Identify Gene Expression Designs With Reduced Burden

**DOI:** 10.1101/013110

**Authors:** Francesca Ceroni, Rhys Algar, Guy-Bart Stan, Tom Ellis

**Author notes:** Please address all correspondence to TE.

## Abstract

Heterologous gene expression can be a significant burden to cells, consuming resources and causing decreased growth and stability. We describe here an *in vivo* monitor that tracks *E. coli* capacity changes in real-time and can be used to assay the burden synthetic constructs and their parts impose. By measuring capacity, construct designs with reduced burden can be identified and shown to predictably outperform less efficient designs, despite having equivalent expression outputs.

---

Robust expression of heterologous genes is necessary for many applications in biotechnology and is central to synthetic biology where predictable fine‐tuning of expression is typically desired^1–3^. However, for engineered bacteria all heterologous expression represents an unnatural load, consuming cellular resources usually allocated to replication, repair and native gene expression (Figure 1A). Gene expression burden is a well-known phenomenon characterised by decreased growth rates that can predispose synthetic constructs to evolutionary instability and can unexpectedly alter their behaviour^4–10^. Burden presents a major barrier to predictable and stable engineering of cells, yet it is largely an unquantified phenomenon, inferred in most cases by tracking growth rate decline^5, 6, 11^. Recent research has begun to explore burden, demonstrating how its impact varies between different *E. coli* strains^6, 11^ and showing how expression load can be measured *in vitro* using cell‐free extracts^12^. However, an improved way of quantifying how heterologous gene expression imposes burden *in vivo* has yet to be described, despite the arrival of new models of bacterial growth that outline the importance of expression resources for the cell^13‐15^.

To advance *in vivo* quantification of burden we developed a fluorescence‐based method to measure in real‐ time the gene expression capacity of bacterial genomes. We built integration vectors to insert a ‘capacity monitor’, a synthetic constitutive green fluorescent protein (GFP) expression cassette, into defined genomic loci of commonly‐used *E. coli* strains (Figure S1); reasoning that because this cassette lacks regulation, changes in GFP expression due to global expression changes will reflect changes in resource availability^16^. To demonstrate how the capacity monitor improves on using growth rates to assess burden, we measured GFP expression rates from the genome of DH10B *E. coli* hosting an operon-expressing plasmid induced at different time‐points during exponential growth (**Figure 1B**). Capacity (determined as GFP production rate per cell) decreases significantly compared to uninduced cells within 30 minutes of construct induction, and this rapid change contrasts with the smaller, slower decreases in growth observed when culture optical density is measured. The fact that capacity changes precede growth rate changes supports the view that decreased expression resources causes growth rate decline and underlines the value of this approach.

**Figure 1.**
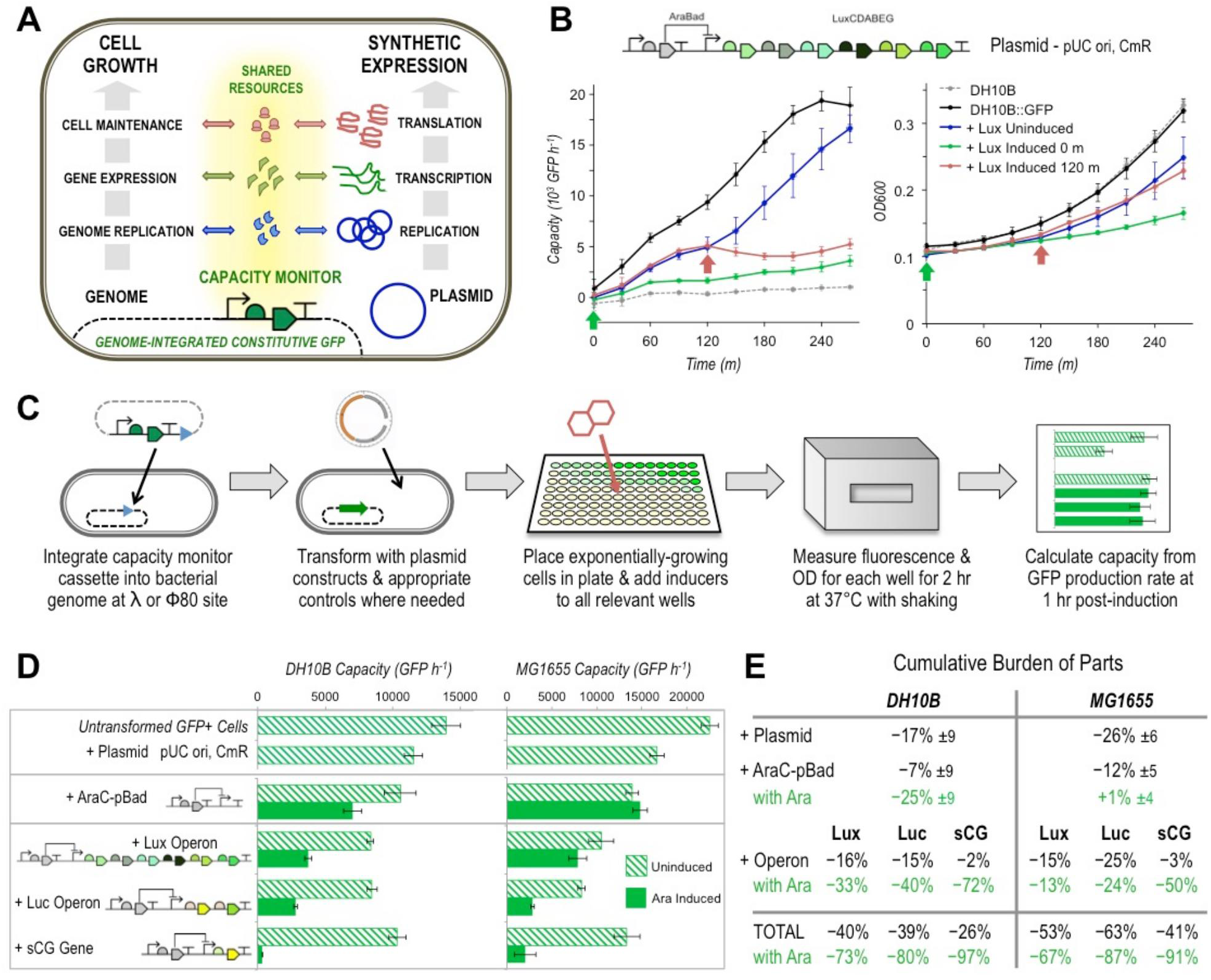
The capacity available for*E. coli* gene expression can be indirectly measured to quantify burden. Schematic representation of the capacity monitor implemented in *E. coli* by genomic integration of a synthetic GFP expression cassette. Changes in the monitor expression rate report real-time changes in the shared resources of the cell that heterologous expression draws upon. **(B)** Effect on the measured capacity (left) and growth (right) of DH10B *E. coli* when expression of a 6-gene *Vibrio* luciferase (Lux) operon from a high-copy plasmid is triggered by L-arabinose induction at two independent time points (0 and 120 min, green and red arrows, respectively) during 4.5 hour of exponential growth. DH10B: GFP have the capacity monitor integrated into the genome at the λ locus, DH10B is the unmodified parent strain. Capacity is determined by the change in total GFP (a.u.) per cell over 1 hour and growth is measured by optical density at 600 nm (OD600). Error bars represent the standard deviation of 3 technical repeats. **(C)** Illustration of method to assay plasmid construct burden in bacterial cells by measuring capacity changes via GFP production. **(D)** The measured capacity of GFP^+^ DH10B and MG1655 cells and cells transformed with plasmid either containing no insert, containing the AraC-pBad regulator-promoter cassette, or containing complete constructs giving Lux operon expression, firefly luciferase operon expression (Luc), or expression of the synthetic human gonadotropin protein (sCG). Solid green bars indicate L-arabinose-induced samples 1 hour post-induction and hatched bars indicate uninduced. Error bars represent the standard error of 3 independent repeats. **(E)** The loss of capacity due to the parts, plasmid and inducers used for the constructs shown in **D** in DH10B and MG1655 cells. Cumulative burden is expressed as a percentage of the total capacity measured for untransformed GFP+ cells, with ±% showing the standard deviation. Calculation of values is described in the Methods.

Having demonstrated that capacity changes within an hour of induction can be used to measure the burden a construct imposes on its host, we developed a routine plate-based assay to measure the performance of synthetic constructs in DH10B and MG1655 *E. coli* (Figure 1C). This can be used to quantify the impact of induced and uninduced constructs and dissect how construct composition causes burden by comparing the capacity of empty plasmids to those containing construct parts (Figure 1D). In both DH10B and MG1655 cells, capacity is decreased by plasmid maintenance and further decreased as the DNA parts that compose constructs are added. Capacity decreases in both strains lead to subsequent growth rate decreases of similar magnitude (Figure S2). Induction of expression from constructs gives the greatest decreases in capacity and these vary significantly depending on the size and function of the genes being expressed. The burden caused by the different parts of expression constructs can be calculated from measured capacity and used to identify those with the greatest cost to the cell (Figure 1E). As well as determining the cost of the different parts this also reveals that the impact of these can vary between different strains. The induction of plasmid-encoded AraC was seen to be a cost to DH10B cells but not to MG1655, presumably due to the different genotypes of these strains (DH10B has deletion of multiple Ara genes). The assay also allows interchangeable parts to be assessed so that different design options can be compared. AraC-based regulation imparts less burden than other well-known inducible systems such as those employing the TetR, LuxR and LacI transcriptional regulators (Figure S3). RNA-based regulation, for example with an inducible riboswitch, has even less cost to the cell due to less resources being required for its operation(Figure S4).

The magnitude of burden imposed by heterologous expression is a product of how much capacity its expression uses and whether the proteins expressed have specific functions that are detrimental to the cell. While redesigning constructs to not produce proteins that interact negatively within the host is a challenge specific for each case, we reasoned that it would be possible and generalisable to redesign constructs to have reduced burden by having less impact on expression capacity. To explore this, we used the capacity monitor to assess a library of designs of inducible constructs that express a large protein that has no function in our cells. For this we chose the*C. violaceum* enzyme VioB, as neither its substrate nor product are present in *E. coli*. This was fused to a downstream mCherry reporter to enable quantification. Using parts and design tools developed to enable predictable tuning of gene expression, we varied the VioB-mCherry output within the library by changing plasmid copy-number, the core promoter and ribosome binding site (RBS) strengths and using *E. coli* codon-optimised and de-optimised versions of the VioB coding sequence (See Figure S5 and **Methods** for full details). Using the capacity monitor assay, we quantified the burden placed on DH10B cells whilst simultaneously measuring construct output as the rate of red fluorescence production per cell (Figure 2A). The same analysis was also performed for MG1655 cells (Figure S6). In both cases all constructs displayed low output with high capacity and growth when uninduced, except construct H1, which had leaky promoter repression. A variety of different outputs were measured 1 hour post-induction, which verified the construct changes designed to increase or reduce expression. Generally greater output was matched by decreased capacity and the measured capacity correlated strongly with the growth rate (Figure 2B). In contrast, a precise inverse correlation between construct output and cell capacity was not observed (Figure 2C), and revealed instead that some designs proportionally take up greater capacity per construct output (*e.g.* construct M1) and are therefore less *efficient* for the host cell; where efficiency defines a measure of simultaneously maximising both construct output and cell capacity.

**Figure 2.**
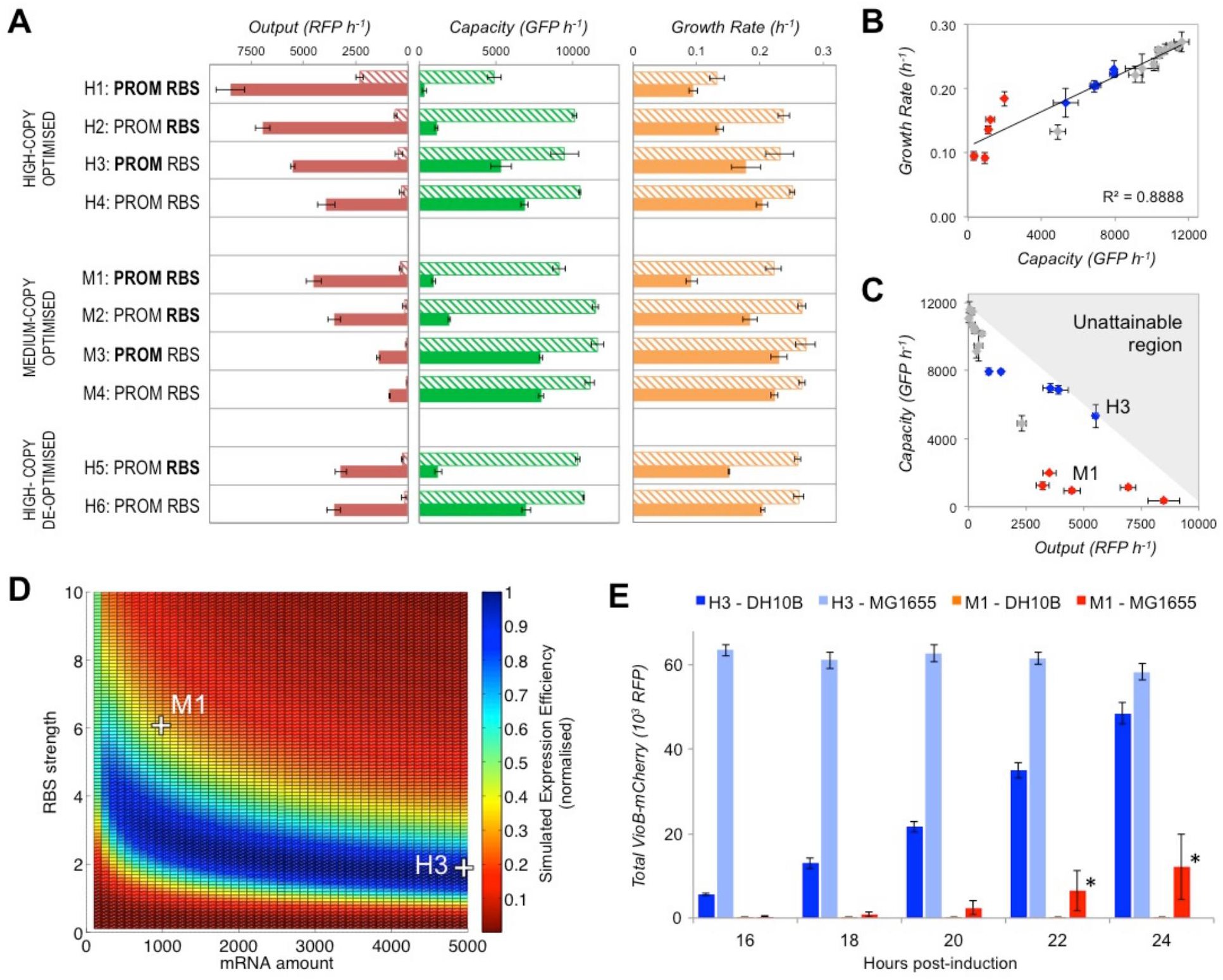
Exploring the relationship between capacity, growth rate and construct output for a combinatorial library of constructs that inducibly-express a VioB*-*mCherry fusion protein. **(A)** Measurement of construct output, capacity and growth rate of different designs 1 hour post‐induction in DH10B cells in exponential growth. Bold text indicates the use of strong core promoter and RBS sequences, and regular text indicates weaker versions. Solid bars show L‐arabinose-induced samples and hatched uninduced. Capacity is measured as before and Output is measured as the change in total red fluorescent protein (RFP) detection per cell over 1 hour. Error bars represent the standard error of 3 independent repeats. **(B)** Plot of growth rate versus capacity for the library constructs. Grey data points are uninduced constructs, blue are induced weak RBS constructs and red are induced strong RBS constructs. Correlation coefficient (R-squared value) is determined by linear regression analysis. **(C)** Plot of capacity versus output for the same constructs. Constructs H3 and M1 and an unattainable region beyond total capacity (grey shading) are highlighted. **(D)** Heatmap of simulated expression efficiency when mRNA amount and RBS strengths are varied for codon‐ optimised library constructs. Locations of simulations that map to constructs H3 and M1 are shown. Normalised expression efficiency is calculated as the product of output and number of free ribosomes, normalised to the maximum efficiency value. **(E)** Total VioB‐mCherry output of H3 and M1 in DH10B and MG1655 *E. coli* when grown induced for 24 hour in shake-flasks. Output is the total red fluorescence per 200 μl of culture. Error bars show standard error of 3 independent repeat experiments. Large error (*) is seen for M1-MG1655 cultures due to null output escape mutants arising in one repeat.

For our construct library all of the inefficient designs were seen to be those expressing with a strong RBS (Figure 2C, red points). To verify that RBS strength was the cause of inefficiency, we also characterised constructs with an alternative 5’UTR strong RBS sequence and this gave equivalent results (Figure S7). The significance of the RBS in determining burden supports previous work that has highlighted the cell's translational resources, particularly the ‘free ribosome pool’ as the critical factor for optimal cell growth and gene expression^14, 15, 17^. To mathematically test this hypothesis and to guide the design of efficient gene expression constructs, we implemented a translation model of heterologous expression that takes into account ribosome availability^18^. This model describes the steps of translation initiation, elongation and ribosome release along a translating mRNA when available ribosomes are finite. This model can be used to simulate *in vivo* gene expression via a Python script provided at the end of this preprint. Using this model, the impact of varying the amount of mRNA (adjustable by promoter strength and/or plasmid copy-number), the RBS strength and the translation rate (adjustable by codon choice) can be explored. Despite the model only accounting for effects on shared translational resources, it is able to predict the burden phenotypes that we observe experimentally where construct output and capacity monitor expression change as construct design is modified (Figure S8). The model also describes how rare codons or other sequences that slow translation elongation only impact designs with a strong RBS (Figure S9) and this qualitatively matches the experimental data collected for de-optimised *VioB* (Figure 2A). Importantly, the model allows the available design-space for a construct to be simulated to determine the designs with the greatest simulated expression efficiency; those with the greatest construct output for the least use of ribosomes (Figure 2D & **S10**). Both the capacity assay (Figure 2A) and these simulated data predict that a high-copy weak RBS construct (H3) is more efficient than a medium-copy strong RBS construct (M1) with similar output levels, despite the former requiring more construct DNA and mRNA per cell. Greater efficiency should result in faster, more robust growth of cells and so greater construct output from batch growth. To verify this prediction, we scaled-up the expression of H3 and M1 constructs to shake-flask scale and measured total output after overnight induction. In both DH10B and MG1655 cells, total output was significantly greater at any time-point when H3 was used instead of M1. This was largely due to faster growth of H3-expressing cultures (Figure S11) but also due to escape mutations occurring in M1 cultures that led to the growth of cells with no construct output (Figures S12 & S13). These results demonstrate that expression designs with reduced burden give greater yields and superior genetic stability, and that selection of such constructs can be achieved by measuring capacity and accounting for its importance in designs. Importantly, the underlying model relies on generic assumptions that typically hold true in various contexts^18^. Consequently this model can aid in forward design by predicting the burden of expressing alternative genes. As a test of its predictive use in a different context, we employed the same model to correctly predict the effects on cell capacity and construct output of expressing an unrelated short protein (scFv) both with and without codon-optimisation Figure S14).

Quantifying capacity as described here offers a practical route to measuring the burden of heterologous expression in real-time and provides vital new information to direct efficient engineering for synthetic biology. Loss of capacity is detected rapidly upon induction of gene expression and precedes and presumably causes growth rate decreases as expression resources become depleted (Figures 1B & S2). Where the genes expressed have no specific function deleterious for cells, measured capacity strongly correlates to growth rate (Figure 2B), meaning that it can predict the growth rate of engineered cells and determine which constructs will perform better in long-term culturing. Using the capacity monitor, the impact of different parts and designs can be assessed and these can then be optimised to reduce burden or increase it if desired. Specifically in the case of expression of a large protein, design changes that increase translational efficiency, such as using a weaker RBS and optimised coding sequence, reduce the burden on cells by preventing ribosomes getting queued on translating mRNAs where they are unnecessarily sequestered from the free ribosome pool. Avoiding ‘over-initiation’ of translation is emerging as an important new design rule^19^ and our model suggests that the optimal RBS is one that recruits ribosomes into elongation at a rate identical to (or just below) the rate at which they move along the slowest section of mRNA.

In cases where two different constructs perform similar tasks, our work suggests that the design with the least impact on the amount of free ribosomes is likely to impart the least burden. In this regard, RNA-based regulators^20^ should be preferable to transcription factors and biological analog circuits^21^ would outperform equivalent circuits made from transcription factor logic gates^22^. Removing the need for continual protein expression from constructs would also reduce burden during long-term culturing. Recently described constructs that write the memory of events into genomic DNA^23^ could be assayed to determine if they give less burden than established bistable switches that require continual transcription factor expression to maintain memory^24^.

By measuring capacity in alternative strains, genotype-specific differences in burden can also be revealed that aid in selecting the optimal host for heterologous expression. Despite DH10B having an inactivated stringent response, its response to induced expression was equivalent to MG1655, indicating the initial burden response to be ppGpp-independent. We believe that the main burden of gene expression is simply a result of gene expression resource depletion, which has the greatest impact on cells growing close to their maximum rate. Since such depletion constitutes a fundamental process common to all dividing cells, it is likely that this work could be extended beyond *E. coli* and be applied to other rapidly dividing organisms, such as *S. cerevisiae* yeast. Already, the vectors we provide for capacity monitor integration can be used in other bacteria with compatible *att* loci^25^ and the accompanying Python script for translation simulation is general and so appropriate for broad use.

## Methods

### Bacterial strains and DNA constructs

Strains MG1655 [K-12 F^-^ λ^‐^ rph-1] and DH10B [K-12 F^-^ λ^‐^ araD139 Δ(araA-leu)7697 Δ(lac)X74 galE15 galK16 galU hsdR2 relA rpsL150(strR) spoT1 deoR φ80dlacZΔM15 endA1 nupG recA1 e14^-^ mcrA Δ(mrr hsdRMS mcrBC)] were obtained from the National BioResource Project (NBRP) Japan. All synthetic genes were codon-optimised for efficient expression in E. coli by DNA2.0 (Menlo Park, CA). RBS sequences were designed using RBS Calculator Software provided by Denovo DNA (State College, PE). The capacity monitor cassette consists of a synthetic strong constitutive promoter, synthetic RBS, codon-optimised superfolder GFP coding sequence and synthetic unnatural bidirectional terminator (see Figure S1 and **Table S1**). To create genomic integration cassettes for this, it was cloned into into CRIM integration vector plasmids pAH63 and pAH153^25^ between existing EcoRI and PstI restriction sites, propagating these in pir-116 electrocompetent E. coli cells (Cambio, UK). These plasmids can then integrate the capacity monitor cassette by CRIM integration into the lambda (pAH63) or phi80 (pAH153) loci of E. coli genomes, co-transforming with helper plasmid pINT-ts^25^ and selecting with 50 μg/ml kanamycin (Figure S1). Subsequent capacity monitor-containing strains (all lambda-integrated in this study) can be propagated without selection and transformed with construct plasmids for testing by electroporation. Details of the design, acquisition and construction of all of control and construct plasmids assessed with the capacity monitor in this study are provided in below.

### Characterised expression constructs

The three expression constructs giving inducible heterologous gene expression are illustrated in **Table S2** and were all obtained via the iGEM Parts Registry (http://parts.igem.org/Catalog) on the high-copy, chloramphenicol-selectable pSB1C3 plasmid. The Lux operon (BBa_K325909: AraC-pBad-LuxCDABEG) and Luc operon (BBa_K325219: AraC-pBad-LRE-Luc) encode L-arabinose inducible expression of 6 luciferase-related *Vibrio fischeri* genes and 2 luciferase‐related *Luciolacruciate* genes, respectively. In both cases all operon genes were codon-optimised for *E. coli* by DNA2.0 (Menlo Park, CA) for the Cambridge 2010 iGEM team. The sCG operon (AraC-pBad-sCG) was constructed by replacing the Luc operon genes in the aforementioned construct with a synthetic gene encoding the human chorionic gonadotropin beta subunit that had been codon-optimised for expression in bacteria by the Virginia 2012 iGEM team. The regulator-promoter-only control construct (AraC-pBad) was generated by restriction digestion and re-ligation of the Luc construct to remove the operon genes. Further controls were the high-copy, chloramphenicol-selectable pSB1C3 plasmid, the low-copy chloramphenicol-selectable pSB4C5 plasmid and the high-copy, ampicillin-selectable pSB1A2 plasmid^26^.

### Inducible expression constructs

Three alternative inducible expression cassette constructs were prepared for Figure S3 using parts taken from the iGEM Parts Registry (See **Table S4**). The Lux Inverter BioBrick BBa_F2620 encoding pTet-LuxR-pLux on high-copy plasmid pSB1C3 was taken directly as it had been previously constructed and described^27^. To obtain a similar construct for TetR regulation (pLac-TetR-pTet), device BBa_I732913 was amplified by PCR to take all parts from the pLacIq promoter (upstream of TetR) to the pTet promoter (*i.e.* excluding mRFP) and this was placed on high-copy plasmid pSB1A2 plasmid using cloning by Gibson Assembly method^28^. To obtain an equivalent LacI regulated construct (p105-LacI-pLac), standard BioBrick assembly was used place a constitutive promoter (BBa_J23105), the LacI CDS (BBa_C0012) with appropriate RBS (BB_B0032), a transcriptional terminator (BBa_B0015) and the pLac promoter (BBa_R0010) in this order within high-copy plasmid pSB1A2. All constructs were verified by analytical gel digest and DNA sequencing prior to use. To induce the TetR regulator anhydrotetracycline (aTc) was added to culture at 100 ng/ml final concentration. For the LacI regulator induction was with isopropyl β-D-1-thiogalactopyranoside (IPTG) at 1 mM final concentration. For the LuxR regulator, induction was with acyl-homoserine lactone (AHL) at 30 nM final concentration.

### Riboswitch constructs

For Figure S4, constructs containing the theophylline riboswitch (pLac-Riboswitch and pLac-Riboswitch-mRFP) were custom synthesised as GeneStrings by GeneArt (Regensburg, Germany) and cloned into pSB1A2 and sequence-verified (see **Table S5**). The sequence of the pLac promoter and mRFP CDS were kept identical to those in pLac-TetR-pTet (Figure S3), and the riboswitch sequence was as previously described^29^. The pLac-TetR-pTet-mRFP construct was the aforementioned BBa_I732913 device cloned into pSB1A2. All constructs were verified by analytical gel digest and DNA sequencing prior to use. To induce the TetR regulator anhydrotetracycline (aTc) was added to culture at 100 ng/ml final concentration. To induce the riboswitch, theophylline was added to culture at 2 mM final concentration.

### Burden assay and time-course

For the burden assay (Figure 1C), *E. coli* cells with and without plasmid constructs were grown overnight with aeration in a shaking incubator at 37°C in 5 ml defined supplemented M9 media with the appropriate antibiotic. In the morning, 20 μl of each sample was diluted into 1 ml of fresh medium and grown at 37°C with shaking for a further hour (outgrowth). 200 μl of each sample at approximately 0.1 OD600 were then transferred into a 96-well plate (Costar) and placed in a Synergy HT Microplate Reader (BioTek, Winooski, VT) and incubated with 1000 rpm orbital shaking at 37°C for 3 hrs performing measurements of GFP (ex. 485nm; em. 528 nm), RFP (ex. 590nm; em. 645nm) and OD (600 nm) every 30 min. 60 min into the incubation, the plate was briefly removed to add inducer to wells and this time point was set as time zero. All burden assays were repeated independently on three different days and to avoid inhibition of arabinose-induced expression due to catabolite repression, 0.4% fructose was used as the main carbon source. For the burden time-course experiment (Figure 1B), the same procedure was applied, but with the 96-well plate prepared with cells initially at 0.02 OD. The plate was then placed into the Microplate Reader and incubated and measured as above, first with a pre-growth of 60 min to ‘time zero’ before 300 min of growth for the time-course. L‐arabinose was added to appropriate wells at 0 min and 120 min. Final inducer and antibiotic concentrations used in assays were; L-arabinose 0.2%, ampicillin 100 μg/ml, and chloramphenicol 34 μg/ml.

### Data analysis

For the burden assay, growth and protein expression rates per hour were calculated as previously described^5^ with growth rate_t2_ = ln(OD_t3_) - ln(OD_t1_)/ (t3-t1), GFP expression rate_t2_ = ((total GFP_t3_) – (total GFP_t1_)/(t3‐t1))/ OD_t2_, and RFP output rate_t2_ = (((total RFP_t3_) – (total RFP_t1_)/(t3‐t1))/ OD_t2_)+ 400; where t1 = 30 min post-induction, t2 = 60 min post-induction and t3 = 90 min post-induction. 400 was added to all RFP output rates to account for the background red fluorescence of M9 media which decreased at a rate of approximately 400 RFP hr^-1^ as it was consumed by cells during growth. Mean rates and their standard error were determined from the rates measured in experiments done independently on 3 different days. For the burden time-course, the same analysis was applied but calculating values for 30 min time points over 4.5 hours for each sample. Mean OD600 and expression rates and their standard deviation were determined from values calculated from 3 independent wells of the same plate. To calculate the burden cost for each step in the table (Figure 1E), cumulative burden is calculated by subtracting the capacity percentage of the construct from that of the construct immediately above. For Ara induction, the value is subtracted from its uninduced equivalent. For TOTAL cumulative burden (with and without Ara), values are the percentage decrease in capacity of the full system (Lux, Luc or sCG) compared to capacity of untransformed GFP+ cells (100%). Error is expressed as ±% and is calculated from standard deviation values.

### Construction and design of the VioB‐mCherry library

The AraC-pBad-VioB:mCherry construct was generated by Gibson Assembly^28^ of overlapping DNA parts amplified by PCR. Plasmid backbones were amplified from pSB31C3 (high-copy) and from the backbone part of plasmid pLysE (medium copy, Bioline Taunton, MA). The AraC-pBad part was amplified from the aforementioned Luc construct and mCherry was amplified from BioBrick BBa_J06504. The VioB coding sequence was amplified from BioBrick BBa_K274003 and is the second of four *C. violaceum* genes in a pigment-producing pathway that were optimised for *E.coli* expression by DNA2.0 (Menlo Park, CA) for the Cambridge 2009 iGEM team. VioB is a 110 kb chromopyrrolate synthase protein that catalyses the conversion of two molecules of 2-imino-3-(indol-3‐ yl)propanoate into one chromopyrrolate molecule. The MetaCyc database (http://metacyc.org) reveals that neither the substrate nor product for VioB are present in*E. coli* metabolism so the enzyme is predicted to be inactive when expressed on its own^30^. In order to fuse VioB and mCherry together sequence encoding an H-linker peptide^31^ was added to 3’ primers used to amplify VioB and 5’ primers used to amplify mCherry. RBS sequences for the VioB-mCherry CDS were designed using the forward engineering mode of the RBS Calculator^32^ and included into 5’ primers amplifying the VioB region and 3’ primers amplifying the AraC-pBad part. Stronger pBad promoters were generated by incorporating point mutations into the pBad core promoter region as previously described by DTU Denmark 2011 iGEM team **see**( **Table S3**), using inverse PCR of the plasmid with mutation-encoding phosphorylated primers followed by *DpnI* digestion and re-ligation. Incorporation of de-optimised sequence (see below for design explanation) into the VioB-mCherry CDS was done using Circular Polymerase Extension Chain assembly^33^ to replace the distal 267 bp of the VioB CDS (upstream of the H-linker) with a 267 bp gBlock DNA fragment synthesised by IDT DNA (Coralville, IA). All PCR amplifications were done using Phusion High Fidelity Polymerase (NEB, Ipswich MA), all primers were synthesised by IDT DNA (Coralville, IA) and all enzymes used for digestion and ligation were obtained from NEB (Ipswich, MA). All constructs, whether generated in-house or from elsewhere, were verified by analytical gel digest and DNA sequencing prior to use.

### Designing a de-optimised VioB coding sequence

Previous work has highlighted the role that codon optimisation plays in efficient gene expression^34, 35^, however there is no unified view on the mechanisms that lead to reduced translation due to inefficient CDS design. Many support the notion that detrimental sequences in a CDS, either related to codon choice or local mRNA folding, cause ribosomes to pause and create a bottleneck in translation elongation (or ‘ribosome traffic jam’)^36^. Some suggest that rare codons (due to the lack of corresponding charged tRNAs) or rare amino acids are responsible for ribosome pausing in conditions of limited resources^35, 37^; whereas others have instead shown evidence that the presence of anti-Shine‐Dalgarno-like sequences in coding regions directly interact with ribosomal RNA and cause ribosomal pausing^38^. Recent work has shown that ribosomal pausing is not a common occurrence in the transcriptome of exponentially growing *E. coli* ^39^ but this is likely due to the very strong evolutionary pressure to remove such pauses from all highly expressed native genes active during fast growth. Heterologous genes that inadvertently incorporate sequences detrimental to the host cell don’t have the advantage of evolution to remove such pauses and instead codon optimisation is required to achieve this.

To investigate the effects of codon optimisation, we took the optimised version of the VioB‐mCherry construct and re-coded 183 bp close to the 3’end of the VioB coding sequence to contain both rare codons and one instance of the strongest known anti-Shine-Dalgarno-like sequence, AGGAGG. This was done to create a potential translation bottleneck^36, 38, 40^ far away from the 5’ end of the gene where the effect of codon optimisation is disputed^35, 41^ and thus effectively de-optimise the construct. In the 183 bp region all arginine, isoleucine, leucine and proline-encoding codons were exchanged with their equivalent rare *E. coli* codon and this also introduced the AGGAGG anti-Shine-Dalgarno-like sequence^38^. See **Table S3** for details.

### Verifying the VioB-mCherry construct library

Following verification by analytical gel digestion and sequencing of parts, the construct library was verified to determine that design changes led to expected behaviours. To do this we compared the RFP measurements upon induction for all constructs. The mutated stronger pBad promoter gave approximately a 1.5-fold increase in expression above the wild type version as expected. Expression of constructs on high-copy plasmids gave approximately 5-fold greater output compared to equivalents on medium-copy plasmids. Use of the strong RBS gave around 3-fold more output than the weak RBS. The alternative strong RBS gave effectively identical output to the strong RBS. Note that we assumed that outputs from H1 and H2 were close to saturated maximal output and therefore unreliable for assessing how changing a part increases or decreases expression. Approximations of fold-change given here were calculated without considering these constructs. No significant change in output was seen when the inefficient VioB-mCherry CDS was introduced into a construct with weak RBS, but output decreased by 50% when introduced into a strong RBS construct.

### Simulated expression efficiency

Simulation of the experimental results of Figure 2A (Figure S7) and justification of the parameters chosen for this are described below, along with details of the Python simulation script. To determine the efficiency of design space(Figure 2D), simulations of all different mRNA numbers (steps of 100 from 100 to 5000) and RBS strengths (steps of 0.1 from 0.1 to 10) were performed. A heatmap of efficiency at steady state was then generated using MatLab (MathWorks, Natick MA), calculating normalised expression efficiency as the product of simulated construct output and simulated free ribosomes, normalising all calculated expression efficiency values to the maximum value in the simulated design space. Simulated expression efficiency without normalisation was also determined and visualised as heatmaps alongside equivalent heatmaps for the simulated circuit output (Figure S10).

### Simulated experiments

A simulation of our experimental set-up was done by simulating a two-construct system in a way where one construct represents the unregulated gene (the monitor) and the other represents a heterologous construct such as a the VioB‐mcherry construct. Simulation of the model described below allows predictions to be made about changes in the behaviour of a synthetic construct when key control parts are altered, as well as how the expected output from the monitor changes. Description of the simulation code and comparison of simulation results with biological data are given after description of the derivation of the translation model.

To simulate the experimental results, the relative strengths of the parts, as determined in the experimental characterisation above, were used to choose mRNA numbers and RBS strengths for simulations using the translational model that is described below. Other values for the simulations were determined from Bionumbers.org.

As pBad is known to be a strong promoter we assumed each copy of the promoter would produce around 30 copies of mRNA per cell when induced (estimated from Bionumber ID 107667). The stronger mutated version would therefore give 50 copies per cell. The p15A origin medium-copy promoter was set at 20 copies per cell and the pUC origin high‐copy was therefore considered to be at 100 copies per cell. This led to mRNA numbers ranging from 600 per cell for induced M2 and M4 constructs, to 5000 per cell for induced H1 and H3 constructs. Parameters for the simulation of construct expression and capacity monitor expression were: total available ribosomes: 10000, capacity monitor length: 24 (720 bp), capacity monitor mRNAs: 20, capacity monitor RBS strength: 1, capacity monitor elongation rate: 1 throughout, construct length: 100 (3000 bp) for long CDS constructs or 30 (900 bp) for short CDS constructs, construct mRNAs: varying from 100 to 5000, construct RBS strength: varying from 0.1 to 10, optimised construct elongation rate: 1 throughout, de‐optimised elongation rate for long CDS: 1 for steps 1 to 79 and steps 90 to 100 but 0.5 for steps 80 to 89, un‐optimised elongation rate for short CDS: 1 for all steps from 1 to 30, except for 0.5 for steps 6, 14, 20 and 26.

### Shake‐flask scale growth

Constructs H3 and M1 were assessed in DH10B and MG1655*E. coli* over 24 hours of exponential growth in media with inducer, and repeated in full on 3 consecutive days. Starter cultures of H3‐DH10B, H3‐MG1655, M1‐DH10B and M1-MG1655 were grown with shaking at 37°C from individual colonies taken from a plate for 5 hour in 3 ml of supplemented M9 with 0.4% fructose, 34 μg/ml chloramphenicol. Starter culture for each was then diluted to OD600 0.015 and 50 μl of this (~150,000 cells) was used to inoculate batch cultures of 50 ml supplemented M9 with 0.4% fructose, 0.2% L-arabinose, 34 μg/ml chloramphenicol in 500 ml baffled shake-flasks. Shake‐flask cultures were grown with shaking at 37°C for 16 hours, and then 1 ml of culture was removed from each flask every 2 hour until 24 hour. 200 μl of removed culture was measured for optical density (OD600, **Figure S11**) and total VioB-mCherry production (RFP output, Figure 2E) using the Microplate Reader. Red fluorescence and green fluorescence (from the capacity monitor) of individual cells in these cultures was simultaneously determined by flow cytometry with a modified Becton Dickinson FACScan flow cytometer capable of parallel measurement GFP and RFP. Cells were diluted in water and passed through the cytometer for 30 s. A 488-­‐nm laser was used for excitation of green fluorescence detecting through a 530 nm band pass filter (FL1-­‐H). Red fluorescence used a 561 nm laser and 610 nm filter (FL2‐A). Data analysis and presentation (**Figure S12**) was performed in FlowJo (Treestar Inc.), gating forward and side scatter appropriately for *E. coli* cells.

### Plasmid mutation analysis

Samples were grown with induced expression with shaking at 37°C from individual colonies for 24 hour in 15 ml tubes in 5 ml of supplemented M9 with 0.4% fructose, 0.2% L‐ arabinose, 34 μg/ml chloramphenicol and L-arabinose inducer. Samples that had reached OD600 0.6 or above were then screened for red and green fluorescence by flow cytometry, before plasmid DNA was extracted by standard mini-prep procedure. Plasmid DNA was then sequenced from a forward primer extending from the 5’ end of the pBad promoter, giving ~900 bases of sequence data covering the pBad promoter, RBS and start of the VioB coding sequence. Where DNA sequencing failed, analysis of plasmid size was done by gel electrophoresis after restriction digestion.

### scFv Constructs

High-­‐copy plasmids expressing an scFv gene from AraC‐pBad were constructed for **Figures S14, S16 and S17**. Constructs were assembled by purchasing as DNA GBlocks (IDT) four scFv CDS regions with the same strong synthetic RBS sequence (see **Table S6**). These were cloned in place of the VioB‐ mCherry CDS from construct H1 using restriction digestion and ligation to directly swap the RBS‐CDS regions while maintaining the promoter and plasmid regions. Before assaying, all constructs were verified by analytical gel digest and DNA sequencing. Construct F1 consists of AraC‐pBad‐scFV‐A, where scFv‐A consists of a synthetic RBS designed by the RBS Calculator^32^ to be strong (95000 a.u.) followed by the optimised A1 scFv CDS previously described to be codon optimised^34^. Construct F2 is the equivalent of F1 but contains a previously described (A1_24) non‐optimised version of the scFv CDS, which maintains the exact same DNA sequence for the first 45 bp of the CDS^34^. Construct F3 contains the scFv-J CDS designed by the online JCat algorithm^42^ (http://www.jcat.de), which used a codon adaptation index method. Construct F4 contains the scFv‐E CDS designed by Eugene (http://bioinformatics.ua.pt/eugene), a freeware suite using a codon harmonisation algorithm. A further scFV CDS designed by the online Optimizer algorithm^43^ (http://genomes.urv.es/OPTIMIZER) proved difficult to synthesise and could not be cloned into the appropriate plasmid, so was not assessed in this study. For constructs F3 and F4 the sequences for scFv-J and scFv‐E CDS were obtained by optimising the scFv-A CDS downstream of the first 45 bp for*E. coli* K-12. In both cases two codons within the C-terminal HisTag then had to be manually swapped to remove repetitive sequence prohibiting commercial gene synthesis. The firsts 45 bp of the CDS was kept identical for all constructs (F1, F2, F3 ad F4) to ensure that the same strong RBS could be used for all.A weak RBS version of construct F1 (construct F5) was prepared by using site directed mutagenesis to introduce a single base pair mutation into the RBS sequence of construct F1 to change its RBS strength to a predicted strength of 9500 a.u. (calculated by the RBS Calculator in reverse mode^32^) Before assaying, all constructs were verified by analytical gel digest and DNA sequencing prior to use. The burden assay was performed as before.

### Western Blot

To verify and measure scFv production a Western blot (**Figure S16**) was performed and analysed. DH10B *E. coli* containing constructs F1, F2, F3, F4 and F5 were grown overnight at 37°C in shaking incubator in 15 ml tubes in 5 ml of supplemented M9 with 0.4% fructose and 34 μg/ml chloramphenicol. In the morning, each culture was diluted 1:7 into 5 ml of fresh medium with and without induction with0.2% L‐arabinose. After 4 hours of growth, OD600 was measured and 1 ml of cells diluted to OD600 = 0.4 were harvested. Pellets of these cells were re-suspended in 160 μl distilled water mixed with 40 μl 6x SDS‐PAGE loading buffer. Samples were then denatured by boiling at 94°C for 10 minutes. 8 μl of each sample were loaded and separated by 12% SDS-PAGE alongside a positive control sample that verifies the antibody functionality (extract from *P. pastoris* GS115 expressing a known single chain antibody fragment with C‐ terminal HisTag) and 5 μl of the 10‐170 kDa PageRuler Prestained Protein Ladder (Thermo Scientific). Protein was transferred from the gel to membrane (Millipore) using a Trans-Blot SD semi-dry transfer cell (BioRad). Protein detection was performed with an anti-His primary antibody (BioLegends, UK) and the WesternBreeze Chromogenic Western Blot Immunodetection Kit (Invitrogen). Image analysis of the Western blot was performed in ImageJ (NIH) by converting a colour photograph of the membrane into greyscale and measuring the maximum and minimum grey intensity in the region immediately at and around each visual band. Intensity was calculated by subtracting the minimum (local background) from the maximum (band).

## SUPPLEMENTARY NOTE 1: Burden of different expression induction cassettes

Three inducible expression cassette constructs were assessed in Figure S3 as alternatives to the AraC-pBad regulator used throughout this study. These were based on three different commonly used transcriptional regulator cassettes, where a transcription factor regulates a specific promoter via its operator sites. The cassettes tested use three different transcription factors; TetR, which binds to the pTet promoter as a repressor and repression is relieved by addition of anhydrotetracycline (ATc); LuxR which binds to acyl-homoserine lactone (AHL) and then acts as an activator for the pLux promoter; and LacI which binds to the pLac promoter as a repressor and repression is relieved by addition of sopropylI β-D‐1‐ thiogalactopyranoside (IPTG). All three cassettes were expressed from promoters that are constitutive in DH10B and MG1655 cells in standard media, and no gene or operon was placed downstream so that the burden assay only compared the cost ofthe induction system itself.

For all three alternatives, the assay revealed there to be significant burden imparted by the inducer constructs both with and without inducer provided. All three constructs require constitutive expression of their transcription factor, which is a cost that decreases gene expression capacity compared to untransformed cells. In all cases the burden of these alternative constructs is greater than that for the AraC-pBad cassette used elsewhere in this study. The LuxR regulator is particularly costly to DH10B *E. coli*. In both strains the presence of the three inducers does not significantly decrease capacity further, demonstrating that the inducer molecules themselves do not cause any burden. Induction of the LacI construct with IPTG interestingly increases capacity compared to uninduced. A possible explanation is that overexpressed LacI from this construct has detrimental binding to off-target sites in the genome and addition of IPTG relieves this by preventing binding.

## SUPPLEMENTARY NOTE 2: Burden of RNA‐based expression induction systems

To determine whether an RNA-only system can impart less burden than a protein-based one, an inducible riboswitch was compared to a transcriptional regulator system in Figure S4. As there is currently no riboswitch confirmed to work *in vivo* in *E. coli* that responds to the same inducer molecule as a transcriptional regulator, we chose to compare a previously characterised theophylline-activated riboswitch determined to have good on/off characteristics^29^ to the ATc-activated TetR-pTet system used in Figure S3. To ensure a direct comparison of the cost of the system, the same plasmid and upstream promoter was used for both. This required changing the promoter used for the riboswitch from the very strong T5 promoter utilised in previous use of this system^29^ to the weaker pLac promoter.

Significantly less capacity was measured in cells expressing the protein-based regulator (pLac‐TetR‐pTet) compared to those expressing the RNA‐based regulator (pLac-Riboswitch), both when induced and uninduced. Capacity of cells with the RNA‐based regulator was similar to the capacity of cells containing plasmid only, both in MG1655 and DH10B *E. coli*, demonstrating that the riboswitch system alone is an insignificant cost. To verify that the two systems worked as regulators, the coding sequence of the reporter protein mRFP was placed immediately downstream of the constructs and their output was measured by determining RFP production rate as in Figure 2A, while simultaneously measuring burden as before. For both systems, a significant increase in RFP production is observed when induced, confirming that they work as expected. Extending the constructs to include the mRFP sequence gives a minor decrease in capacity, and no significant change in capacity is seen between uninduced and induced samples. Note that for the pLac-Riboswitch-mRFP construct a low rate of RFP production is observed, while a high rate of RFP production is seen for the pLac-Tet‐pTet­mRFP construct. We attribute the difference in construct output to the different mRFP RBS strengths used. The TetR-regulated construct used a strong RBS (BBa_B0034) and the RBS for the riboswitch could not be modified to match this, as the RBS forms a critical part of the riboswitch and cannot be altered without impeding its function^29^.

## SUPPLEMENTARY NOTE 3: Predictive capability of the translation model

To verify the predictive capability of the model and associated numerical simulations beyond VioB-mCherry constructs, we chose to use the model to predict the effects on capacity of expressing a shorter heterologous protein CDS (900 nt, 30 elongation steps, where an elongation step is 10 codons at a time, the approximate width of a ribosome on an mRNA) compared to the large protein CDS previously simulated (3000 nt, 100 elongation steps). Keeping all other parameters the same, the model predicts that decreasing CDS length while maintaining mRNA copy number and RBS strength leads to a significant increase(>3-fold) in capacity of the cell (see **Figure S14A**). To confirm this prediction experimentally we identified a short heterologous protein whose expression from *E. coli* had been previously characterised^34^. The protein selected was a codon-optimised antibody single-chain variable fragment (scFv) fused with a C-terminal HisTag to enable characterisation of expression using Western blot with anti-His antibody staining. The protein has a mass of 32 kDa and a CDS size of 846 bp and was paired with a synthetic RBS sequence predicted to have equivalent strength to the strong RBS for VioB-mCherry was designed using the RBS Calculator^32^. The capacity of DH10B *E. coli* expressing this scFv protein was measured by burden assay using a high-copy construct (F1) of equivalent composition to construct H2 from Figure 2A. Capacity was measured to be significantly higher than previously observed for the equivalent longer VioB construct (**Figure S14B)** and calculated to be approximately 3.5-fold greater, a close match to the simulation predicted increase.

To further explore the predictability of the model we also took the opportunity to compare the effect on capacity with a weaker RBS and with un-optimised codon usage in both simulation and in experiments. For the codon usage, an scFv CDS was available that was previously described to be un-optimised^34^ and this was exchanged with the CDS in construct F1 to create construct F2. For a weaker RBS, we added a point mutation into the strong RBS used in construct F1 which was predicted to result in an RBS strength slightly stronger than that used as the weak RBS for VioB-mCherry in Figure 2A. Before these constructs were experimentally tested, their output and burden were simulated maintaining all other parameters as before. Note that simulating an un‐optimised CDS required a different approach to simulating the de-optimised CDS in Figure S9, which mimicked an intentionally introduced translation bottleneck by placing 10 slow elongation steps towards the CDS end. Instead for an un‐optimised CDS, codon usage was simulated to have slow elongation steps (*i.e.* poor codons) interspersed along the CDS, rather than concentrated at one location.

Simulated data shown in **Figure S14A** predicted that using an un-optimised short CDS would lead to a decrease in construct output and almost no change in cell capacity compared to an optimised CDS. A similar prediction had been made for de‐optimisation of the long CDS equivalent. Likewise, with a weaker RBS, the short CDS simulation predicts half the output of the strong RBS equivalent and much greater capacity, as also predicted before for the long CDS equivalent. Obtaining experimental data to compare against these predictions required both a burden assay to be run for constructs F1, F2 and F5 anda simultaneous Western blot (**Figure S16**) to be performed and analysed to estimate construct output.

The experimental data (**Figure S14B**) for the burden of scFv constructs captures the predicted trend for changing to un-optimised CDS or weaker RBS, but the magnitude of capacity differs from that simulated. The model appears to over‐predict the burden (underestimate the capacity) with strong RBS short CDS (scFv) constructs. This is also observed for long CDS (VioB-mCherry) constructs. For the output of scFv,the experimental data are an excellent match for the data predicted by the simulation, and in this case the model also appears better at capturing the effect of non-optimal codon usage for the short CDS (scFv) constructs, when compared to its ability to predict the output for a similar de-optimised long CDS (VioB-mCherry) construct.

## SUPPLEMENTARY NOTE 4: Assessing codon-optimisation methods

To determine if the burden assay method could be used to assess different codon optimisation algorithms, constructs F1 (scFv‐A, optimised) and F2 (scFv‐B, un-optimised) were compared to equivalent constructs F3 and F4, which respectively contained the scFv CDS sequences generated by by the online JCat optimisation algorithm (scFv-J) and the downloadable Eugene software suite codon harmonisation algorithm (scFv-E). All four were designed to maintain the same DNA sequence as scFv-A for the first 48 bp of the CDS to ensure that the same strong RBS could be used for all constructs (*i.e.* to avoid any alteration in the RBS strength due to local sequence context change). All were expressed on high-copy plasmid with the standard AraC-pBad regulator‐promoter cassette allowing expression to be triggered by induction with L-arabinose.

The burden assay revealed that all four constructs imposed the same capacity cost on DH10B *E. coli,* when uninduced and also had the same magnitude of decreased capacity one hour post‐induction. Likewise, growth rates of all four constructs were comparable. In contrast to this, image intensity analysis of a Western blot (**Figure S16**) after 4 hours of growth revealed different construct outputs for the four constructs when induced, and zero output when uninduced. JCat-optimised scFv (F3) had the greatest output, slightly exceeding that of the previously optimised scFv-A CDS (F1). Eugene-optimised scFv (F4) had equivalent output of the previously described un‐optimised scFv-B CDS (F2).

The results therefore show that the burden assay alone cannot be used to assess different codon optimisation algorithms, as the differences that arise from codon optimisation manifest themselves in the construct output and not in significant changes in either the capacity or growth rate of the cell. This result is actually predicted by our model, which shows that slower elongation and/or bottlenecks during translation lead to less construct output but only decrease capacity by a small amount (see **Figures S9** and **S14**). Assessment of codon optimisation therefore relies on construct output measurement. In the case shown here, JCat is a successful codon optimisation method, whereas Eugene fails to improve scFv production beyond what is seen for an un-optimised version (scFv‐B).

**Figure S1:**
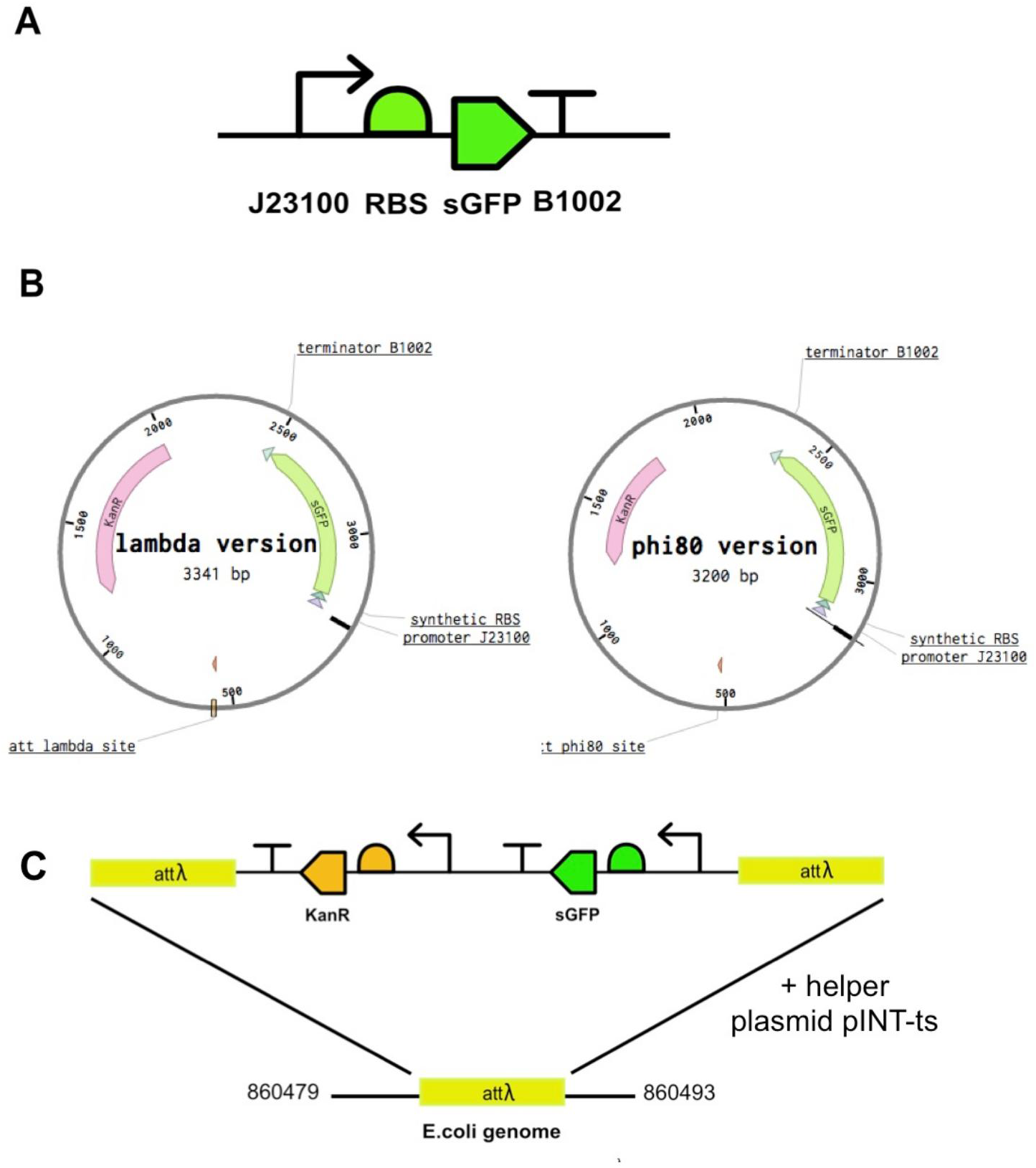
Design of the capacity monitor synthetic cassette and schematic of its integration into the*E. coli*genome. (A) The synthesised capacity monitor cassette consists of a designed synthetic constitutive promoter (BBa_J23100), a designed synthetic RBS, codon-optimised superfolder GFP sequence (sGFP) and a designed synthetic terminator part (BBa_B1002). **(B)** For genomic integration into *E. coli*, the synthesised cassette is inserted into integration plasmids that target either the lambda or phi80 genomic loci. **(C)** The capacity monitor cassette and the downstream KanR selectable marker integrate into the lambda locus of the *E. coli* genome via site-specific recombination with the recombinase provided *in trans* by helper plasmid pINT‐ts.

**Figure S2:**
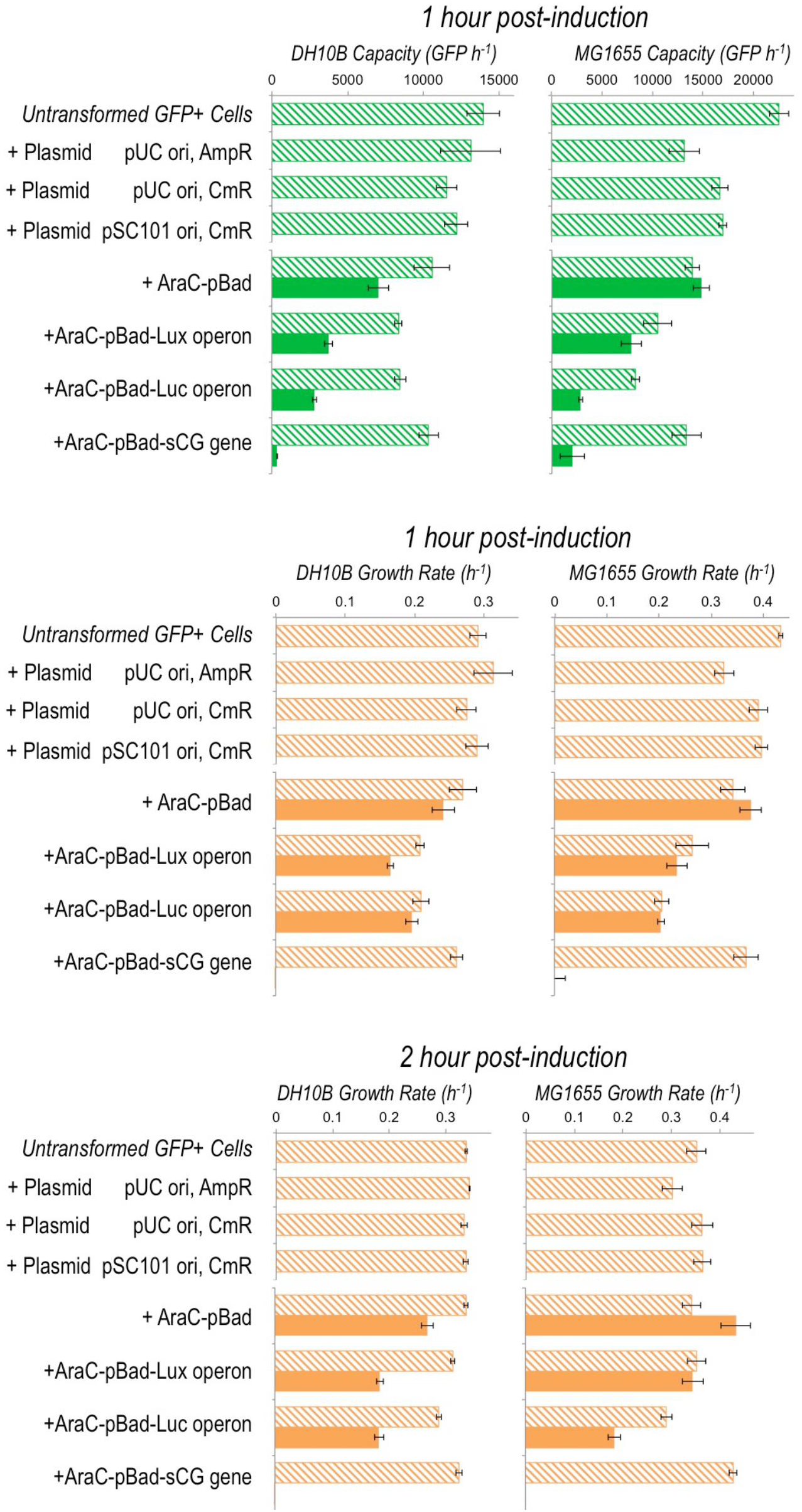
Extended data from Figure 1D showing the burden imposed on MG1655 and DH10B*E. coli*by the differentparts of the Lux, Luc and sCG inducible expression constructs and by three different plasmid backbones. Measured capacity 1 hour post‐induction, and growth rates 1 and 2 hour post-induction are shown. Filled bars indicate L-arabinose induced samples and hatched bars are uninduced. Error bars represent the standard error of 3 biological repeats. Plasmids used here are pSB1A2 (high-copy pUC origin AmpR), pSB1C3 (high‐copy pUC origin CmR), and pSB4C5 (low‐copy pSC101 origin CmR). Constructs are all on pSB1C3 plasmids.

**Figure S3:**
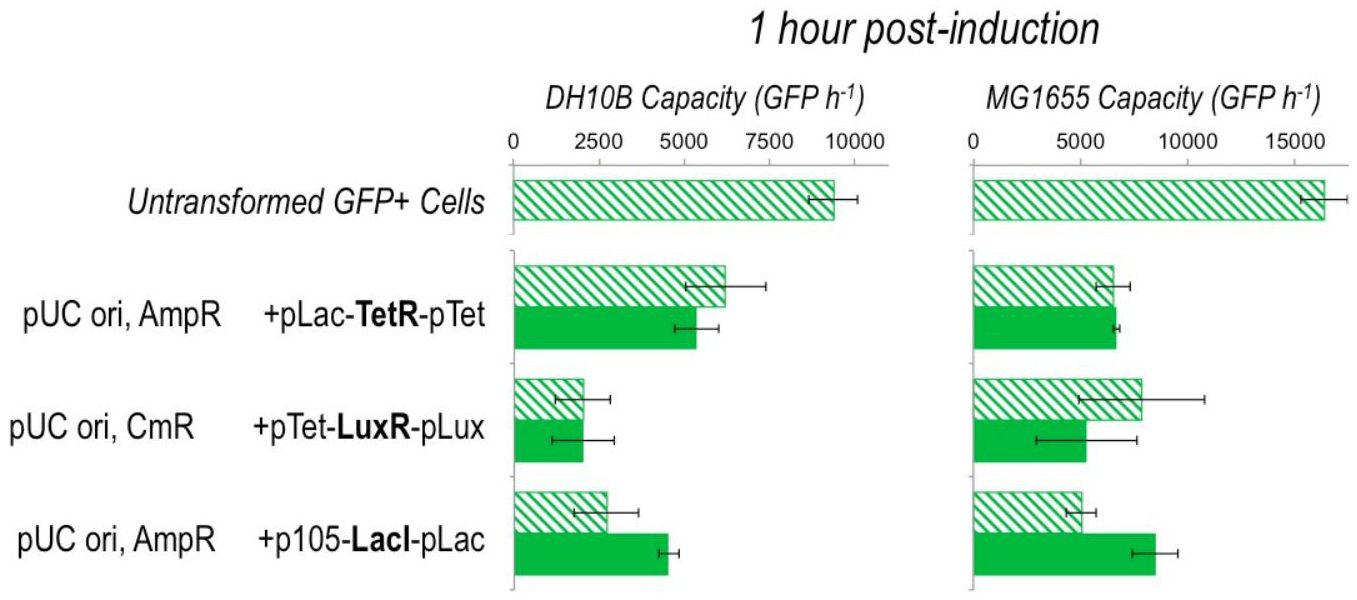
The burden imposed by inducible expression constructs, discussed in **Supplementary Note 1**. The burden ofthe alternatives to AraC‐pBad that give expression following induction was assayed by measuring the effect on capacity and growth of TetR, LuxR and LacI inducer constructs in DH10B and MG1655 cells with the capacity monitor. No gene or operon is placed downstream of the constructs, so measurements give the burden of the induction system with or without inducer only. Graphs show average measured capacity 1 hour post-induction. Hatched bars indicate uninduced and filled bars are induced samples. Error bars represent the standard deviation of 3 independent repeats. Plasmids backbones used are pSB1A2 (high-copy pUC origin AmpR) and pSB1C3 (high-copy pUC origin CmR). Promoters used to express the regulating transcription factors were pLac, pTet and p105 (synthetic promoter BBa_J23105). In their respective constructs these acted as constitutive promoters, with the promoter downstream of the regulator (pTet, pLux and pLac) being regulated.

**Figure S4:**
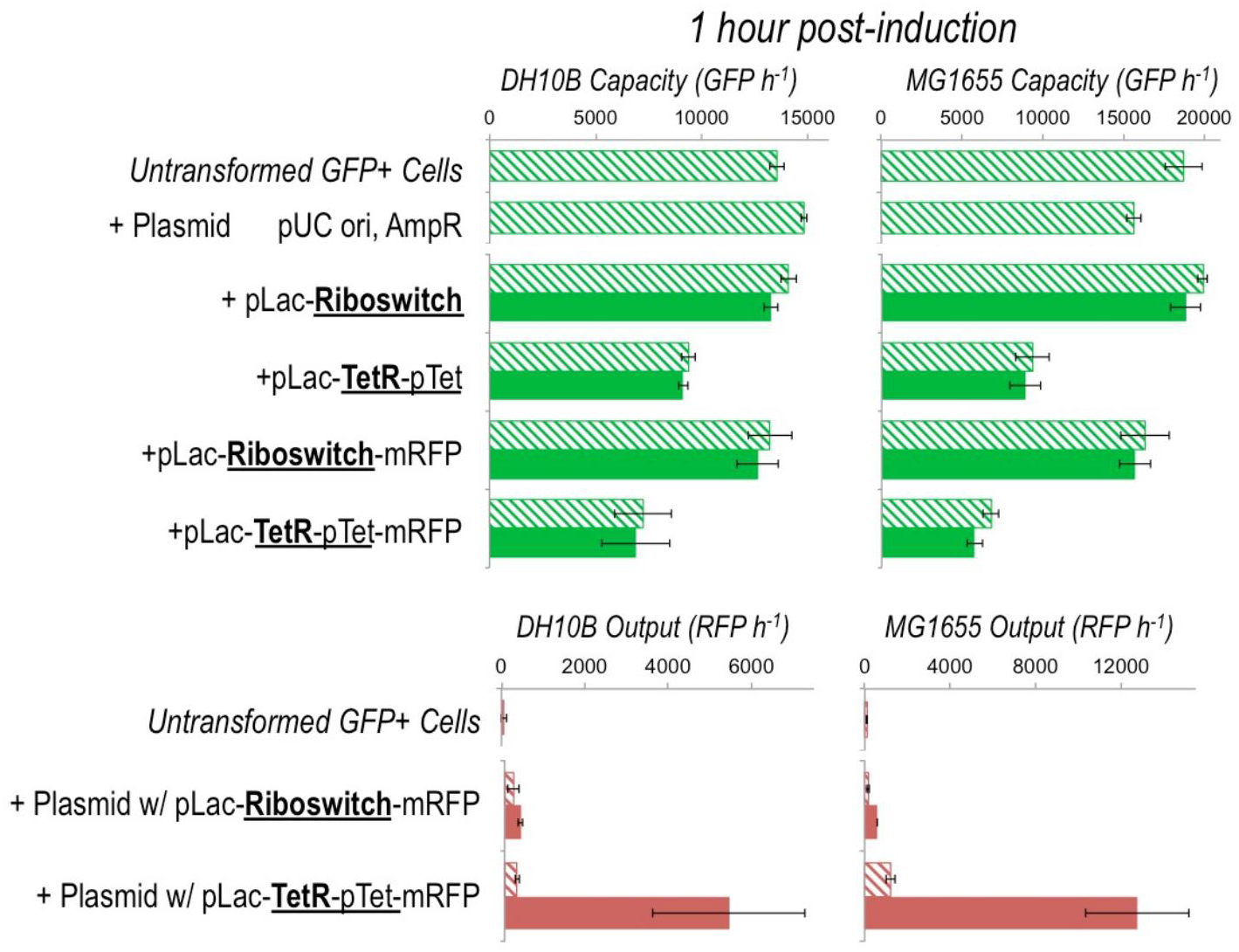
Comparing the burden of RNA-based and protein‐based regulation, discussed in **Supplementary Note 2**.The TetR-pTet inducer system used in Figure S3 was compared to its RNA-only equivalent, a theophylline-­‐activated riboswitch^29^, in DH10B and MG1655 cells with the capacity monitor (top panel). For pLac-Riboswitch and pLac-TetR-pTet, no gene is placed downstream, so measurements give the burden of the induction system with or without inducer only. For pLac-Riboswitch-mRFP and pLac-TetR-pTet-mRFP, the mRFP gene is placed downstream allowing construct output to be determined and to confirm that the induction system works (bottom panel). Graphs show average measured capacity (GFP production rate) and output (RFP production rate) 1 hour post-induction. Hatched bars indicate uninduced and filled bars are induced samples. Error bars represent standard deviation of 3 independent repeats. Plasmid backbone used was pSB1A2 (high-copy pUC origin AmpR). Underlined text highlights functionally equivalent regulation units.

**Figure S5:**
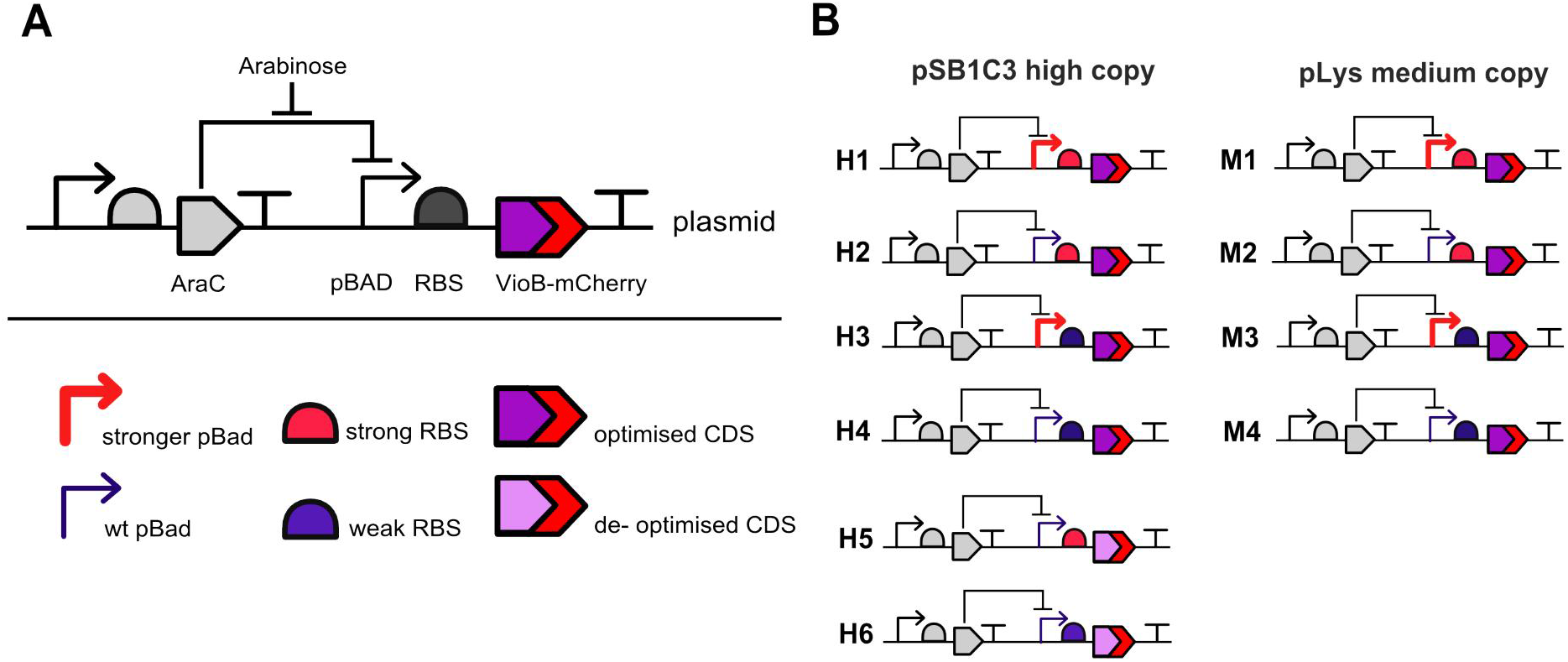
The combinatorial library for different levels of inducible expression of a VioB-mCherry fusion protein. **(A)** Schematic of the construct. In absence of L-arabinose AraC represses VioB-mCherry production by binding to the pBad promoter; when L-arabinose is present it binds to AraC making it an activator of gene expression from pBad. The components of the construct can be changed to tune gene expression. Two versions of the pBad promoter are used, one with wild-type core promoter sequence (*wt pBad*) and one with point mutations that increase output by approximately 1.5-fold (*stronger pBad*). The RBS region was varied using two synthetic 5’-UTR sequences designed by the RBS Calculator^32^ to give *strong* and *weak* translation initiation rates. Codon optimisation is varied by using either an *optimised* version of the VioB-mCherry coding sequence (CDS) synthesised for efficient *E. coli* expression and a de-optimised version where a potential translation bottleneck^36, 40^ is introduced (see Methods) **(B)** Schematic of the library. Construct copy-number was varied using two different chloramphenicol-selected plasmid backbones, one with a *high-copy* pUC19-derived pMB1 origin of replication (pSB1C3) and another with a *medium-copy* p15A replication origin (pLys).

**Figure S6:**
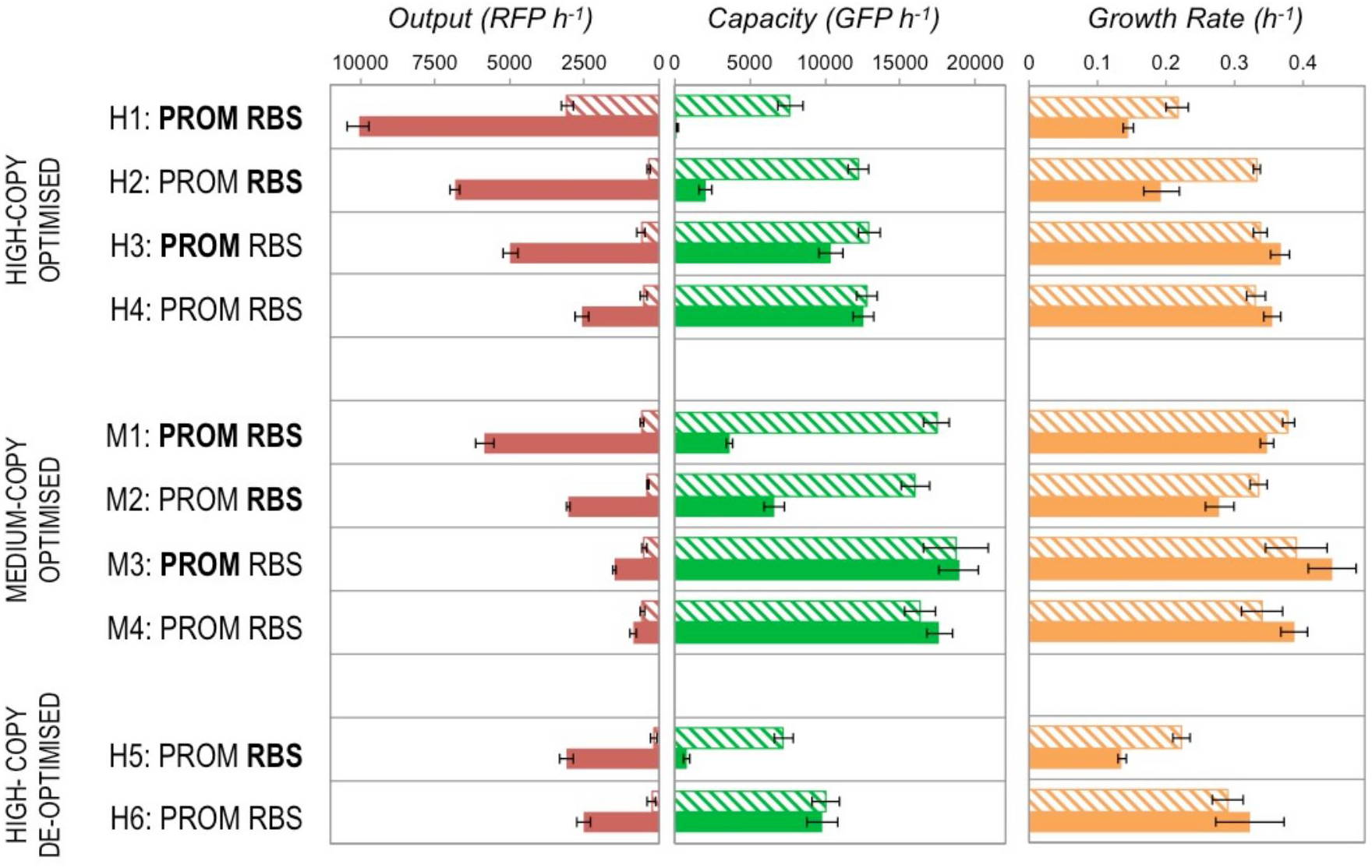
The capacity, growth rate and construct output produced by the VioB-mCherry combinatorial library 1 hourpost-induction in MG1655 cells in exponential growth. Bold text indicates the use of strong core promoter and RBS sequences, and regular text indicates weaker versions. Filled bars indicate induced samples and hatched bars uninduced. Capacity, output and growth rate are measured as for Figure 2A. Error bars represent the standard error of 3 biological repeats.

**Figure S7:**
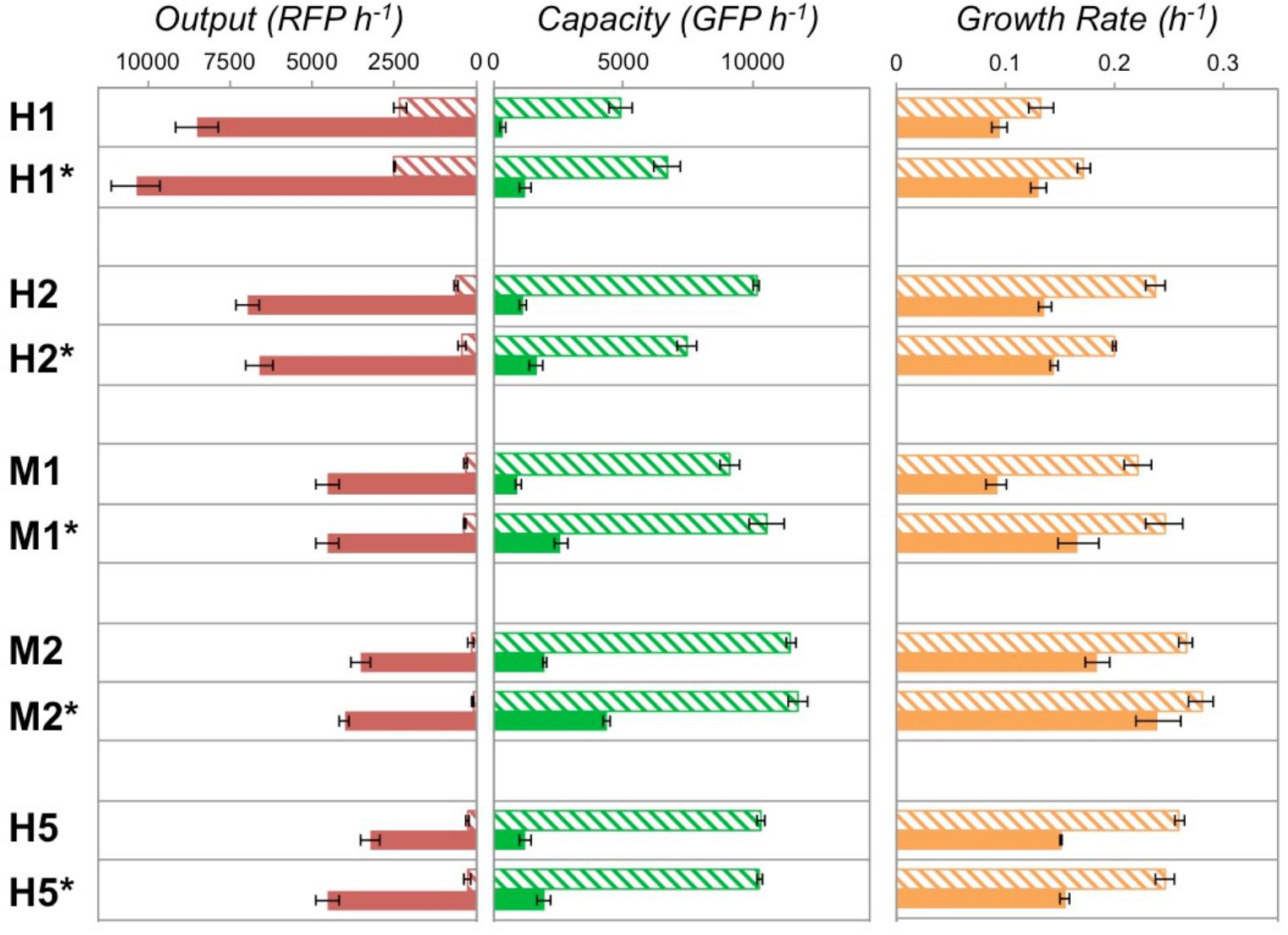
Comparison of construct output and DH10B capacity and growth rates 1 hour post-induction for VioB‐mCherry constructs with strong RBS (H1, H2, M1, M2 and H5) and modified versions of these constructs (H1*, H2*, M1*, M2* and H5*) with a different VioB‐mCherry 5'UTR sequence encoding an alternative strong RBS sequence.

**Figure S8:**
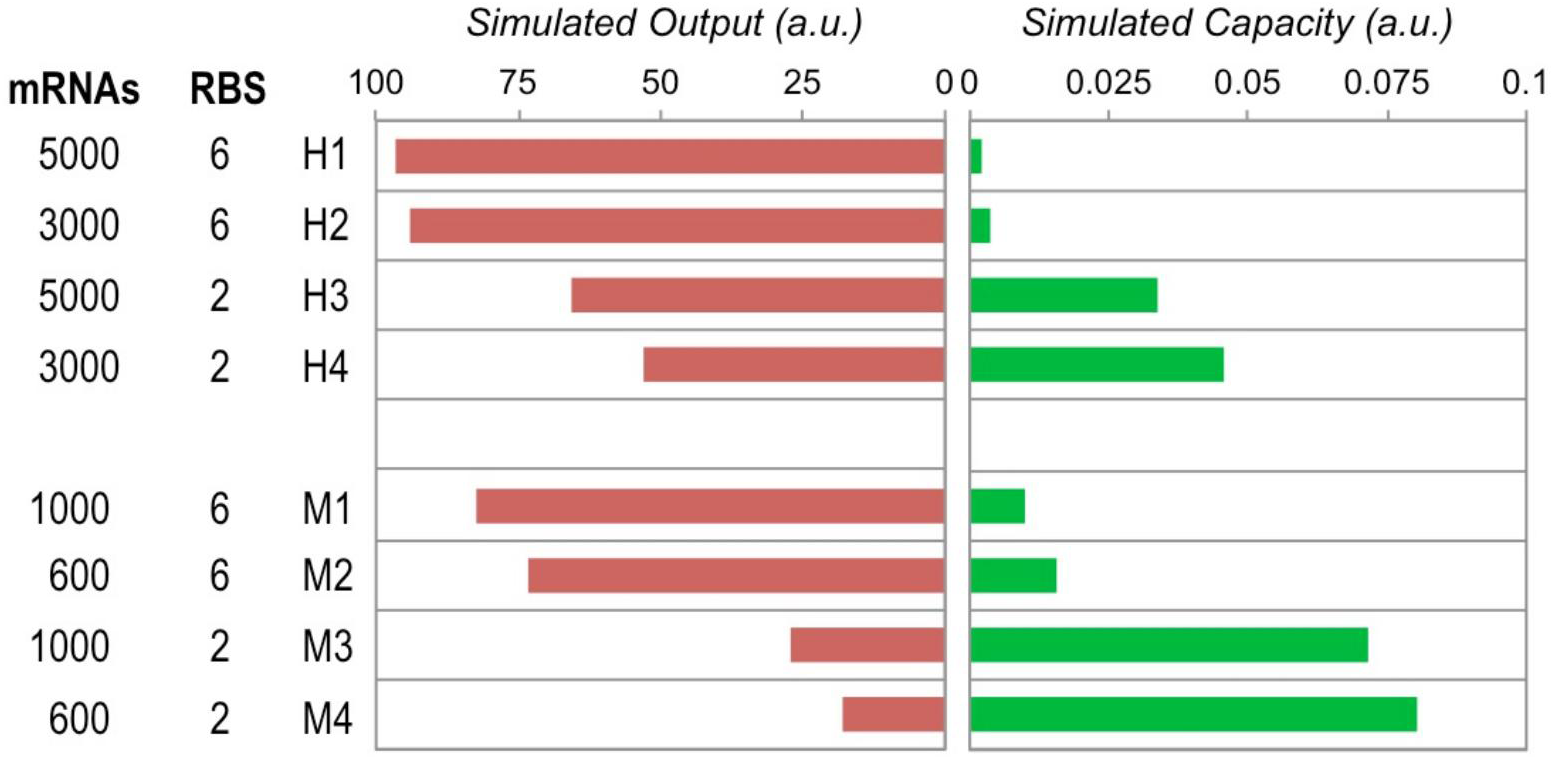
Output and capacity at steady state as determined by a translational resource model for simulated constructs with mRNA numbers and RBS strengths varied to match the relative changes between experimental constructs H1 to H4 and M1 to M4 (Figure 2A).

**Figure S9:**
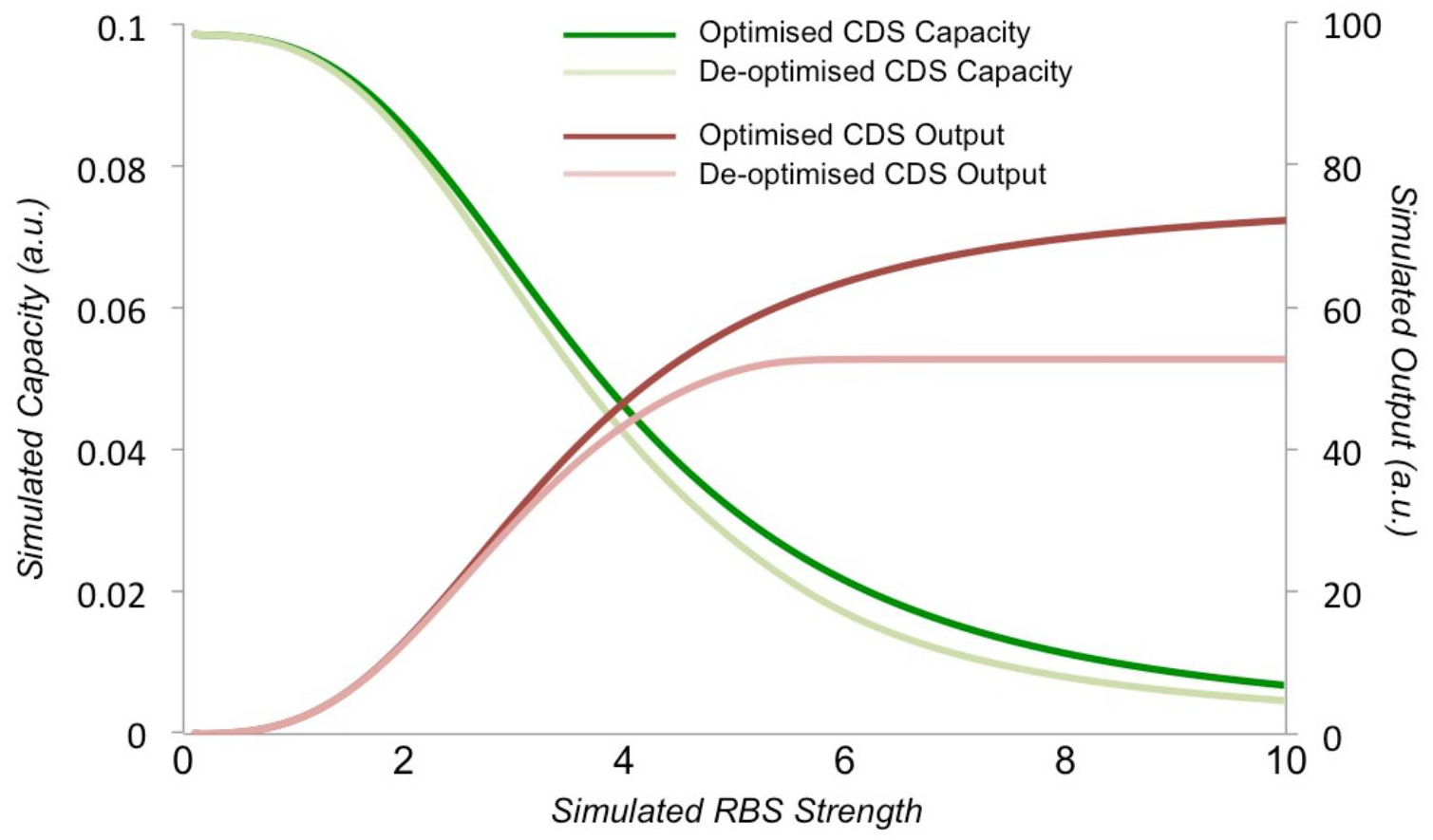
Output and capacity at steady state as determined by a translational resource model for simulated constructs with mRNA numbers and RBS strengths varied to match the relative changes between experimental constructs H1 to H4 and M1 to M4. The simulated effect of varying RBS strengths for two versions of a synthetic construct, one with an optimised coding sequence (CDS) and one with a de‐optimised CDS. At weak RBS strengths CDS optimisation has no effect on output as it is not limiting (see also Figure S10). At strong RBS strengths only the optimised CDS can attain greater output and with a de‐optimised CDS less output is seen for a similar loss in capacity, matching experimental observations in Figure 2A (compare H2 and H5 for strong RBS, and H4 and H6 for weak RBS). The output from the simulated construct is shown in red shades with right axis and the simulated output of a capacity monitor is shown in green shades with left axis. Simulation settings were the same as for Figure 2D, with 400 mRNAs used for the construct. The simulated elongation rate throughout the optimised CDS was set to 1 for all 100 translation elongation steps. For the de‐optimised CDS the simulated translation elongation rate was set to 1 for steps 1 to 79 and 90 to 100 but set to 0.5 for steps 80 to 89, mimicking a translation bottleneck due to rare codons or another sequence‐specific effect.

**Figure S10:**
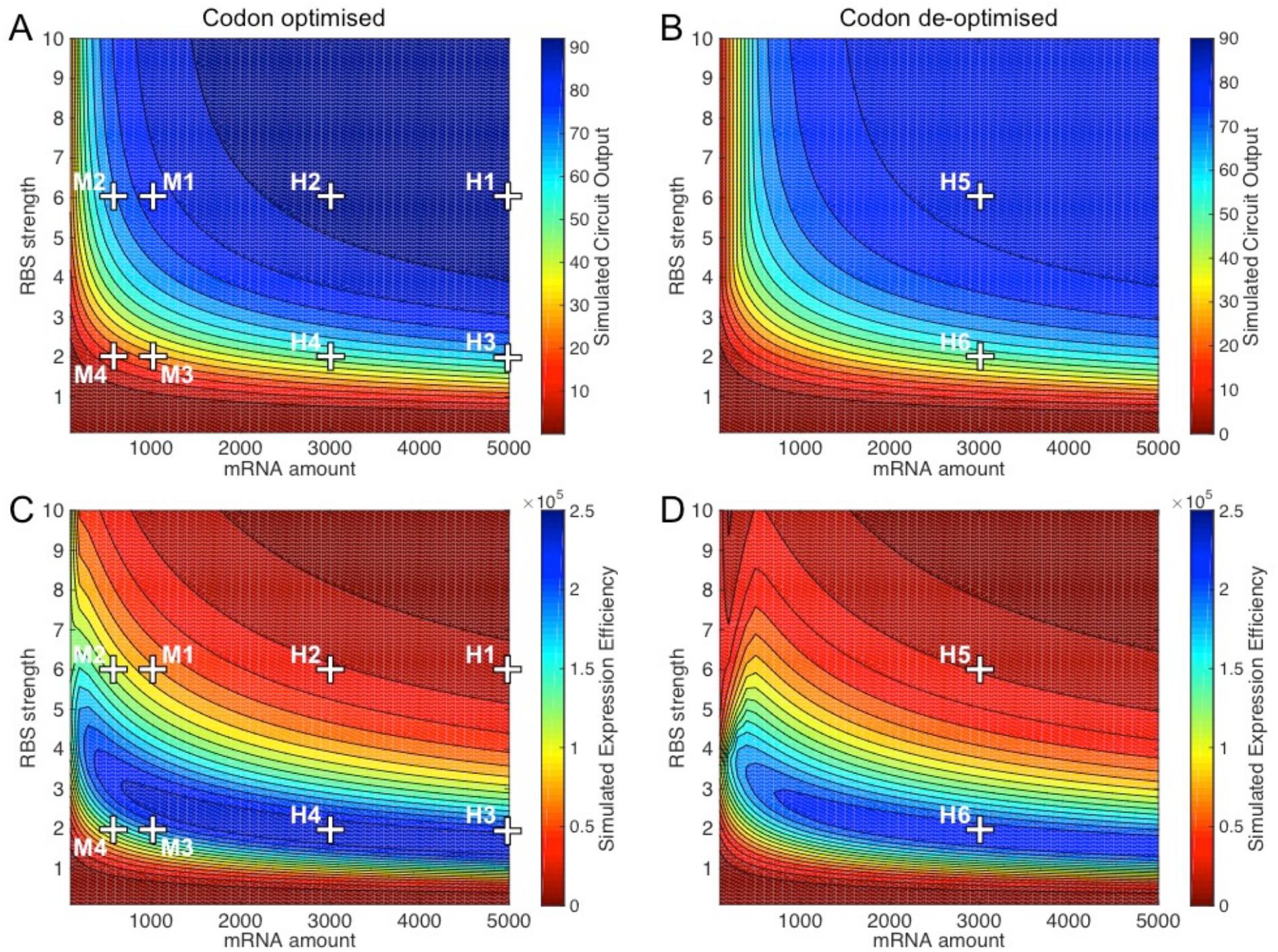
Heatmap of simulated construct outputs for codon optimised (**A**) and de‐optimised **(B**) long CDS constructs alongside heatmaps of simulated expression efficiency for codon optimised (**C**) and de-optimised (**D**) constructs. Locations of simulations mapping to all constructs assayed in Figure 2A are shown. Simulated circuit output is the simulated production rate of the circuit protein at steady state. Simulated expression efficiency is calculated as the product of circuit output and number of free ribosomes available at steady state.

**Figure S11:**
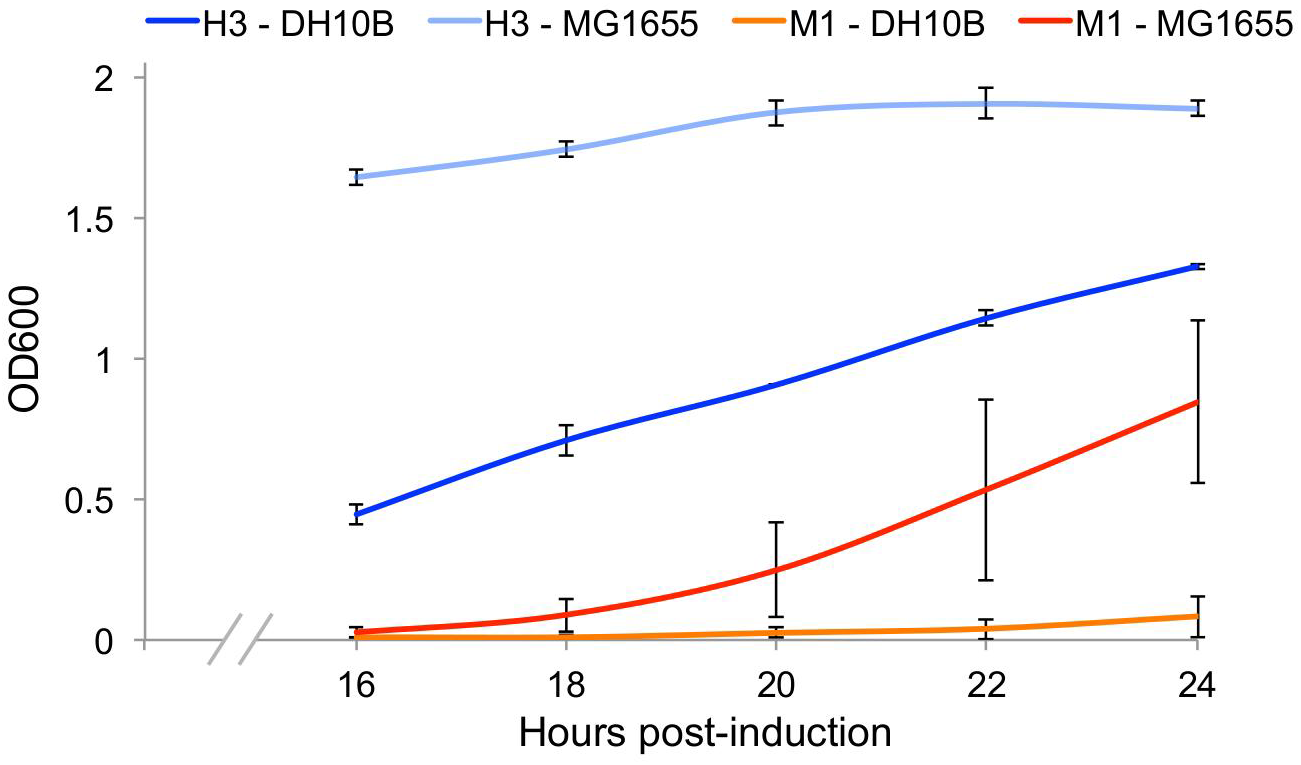
Optical density of DH10B and MG1655*E. coli*hosting the H3 and M1 expression constructs (Figure 2E)when grown under induction conditions for 24 hours in shake-flasks. Growth was performed at 37°C in 50 ml media in 500 ml baffled flasks with orbital shaking. 200 μl of culture was removed every 2 hours between 16 and 24 hour post-induction, and optical density determined by measuring OD600 while simultaneously measuring RFP for Figure 2E. Error bars represent the standard error of 3 independent repeat experiments done on consecutive days.

**Figure S12:**
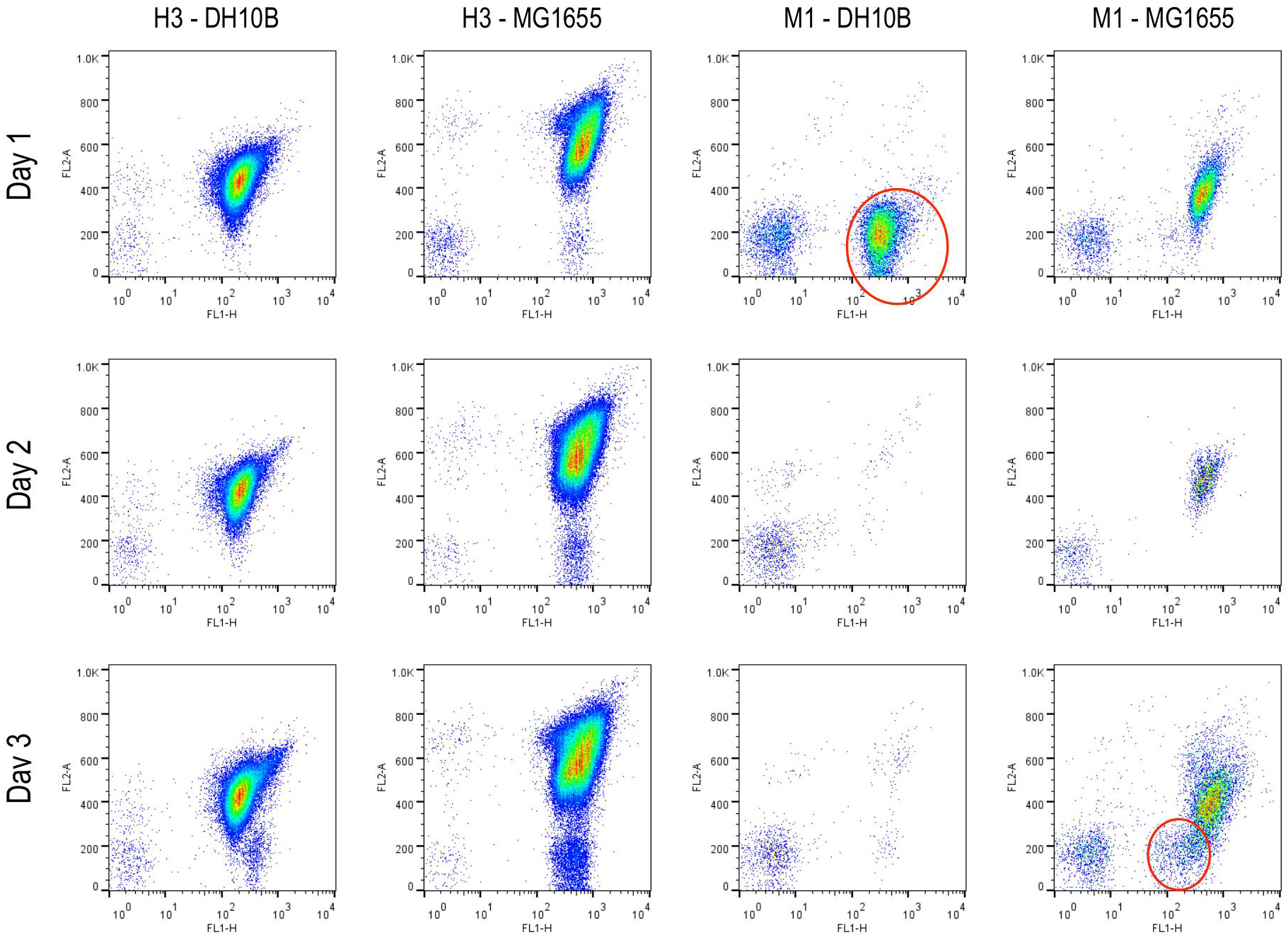
Flow cytometry analysis of the GFP (x‐axis: FL1-H) and VioB-mCherry (RFP, y-axis: FL2-A) content of DH10Band MG1655 *E. coli* hosting the H3 and M1 expression constructs 16 hours post-induction (see Figure 2E and FigureS10). Results are shown for 3 independent experiments done on consecutive days. Populations where escape mutants no longer express VioB-mCherry are highlighted in red.

**Figure S13:**
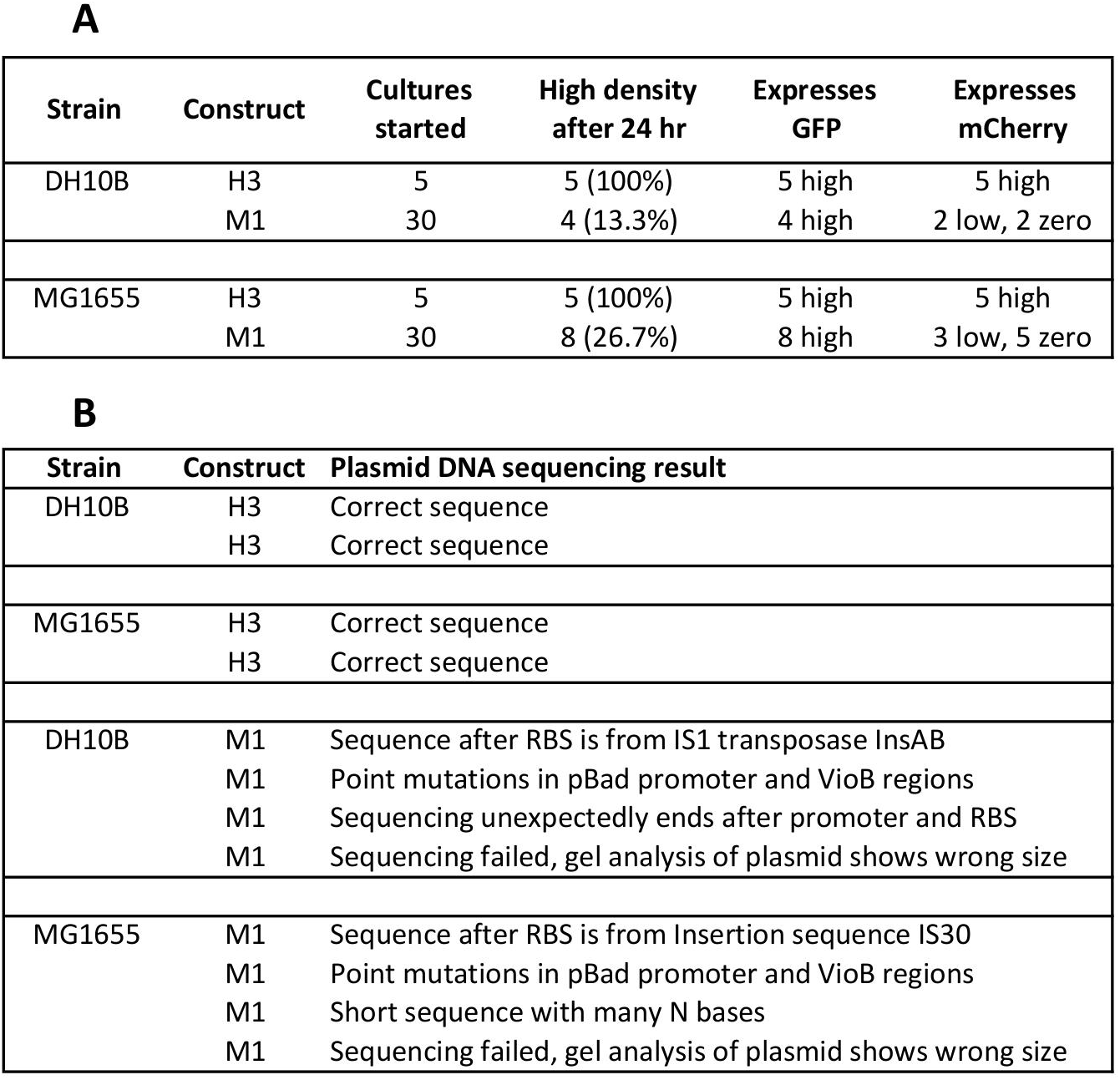
Analysis of escape mutant populations following 24 hour growth in the presence of induction. H3 and M1constructs in capacity‐monitor containing MG1655 and DH10B cells were grown with shaking at 37°C from individual colonies for 24 hour in 5 ml of supplemented M9 with 0.4% fructose, 0.2% L‐arabinose, 34 μg/ml chloramphenicol and L‐arabinose inducer. For H3 in each strain, 5 colonies were grown, and for M1 in each strain 30 colonies were grown. (A) The number of colonies that reached high-density after 24 hours growth and the fluorescence of these. (**B**) For each strain, two H3 and four M1 samples that reached high density were plasmid-prepped and the plasmid sent for DNA sequencing to determine construct integrity.

**Figure S14:**
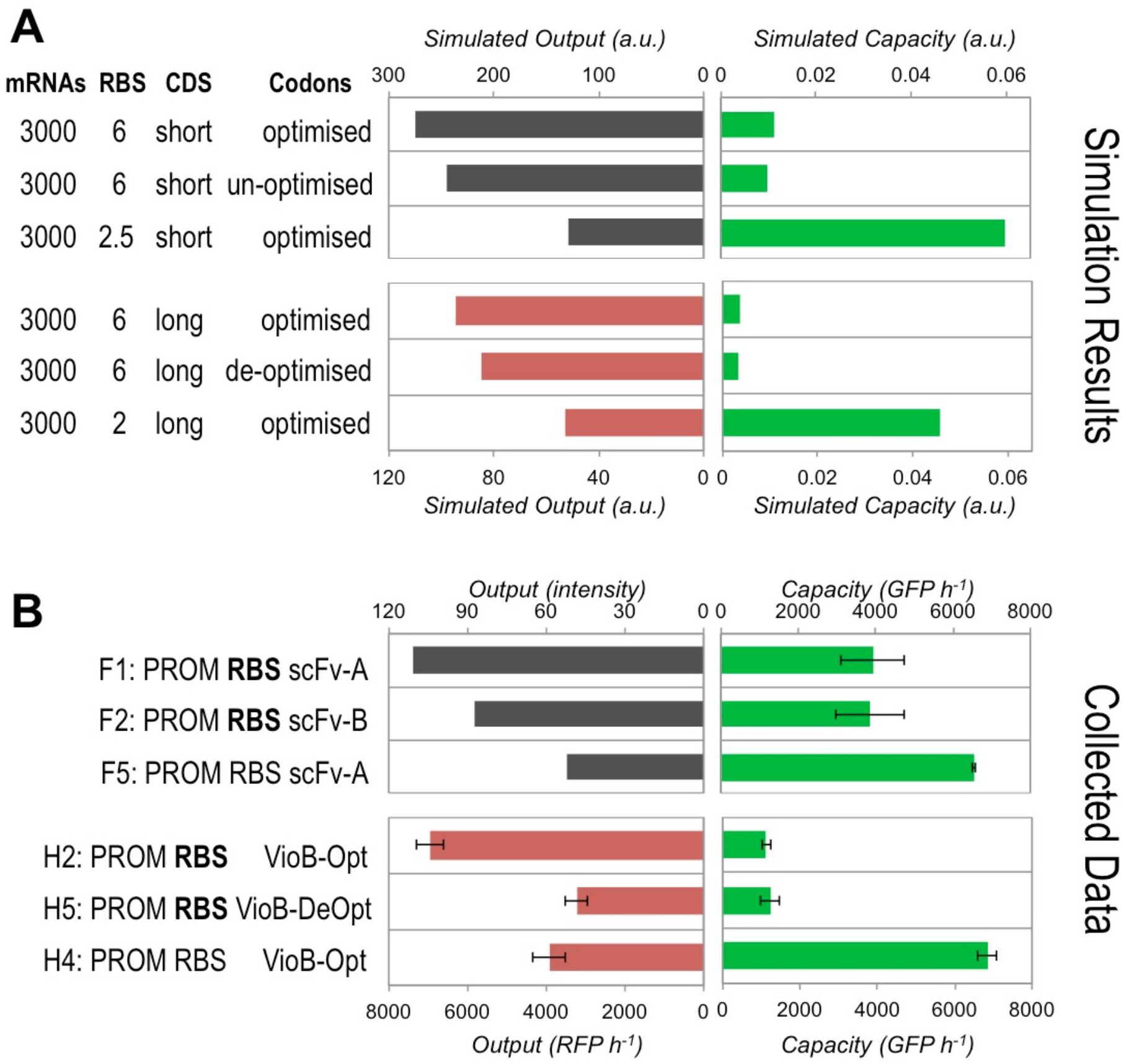
Comparison of simulation predictions and actual obtained results for the capacity and output of scFv constructs expressed in *E. coli* DH10B cells with the capacity monitor. See **Supplementary Note 3** for discussion of these results. **(A)** The model was used to simulate the steady state burden of expressing a coding sequence (CDS) shorter than the long CDS used in Figure 2D. Heatmaps for the design space for short CDS constructs are shown in **Figure S15**. The effect of not having codon optimisation and switching to a weaker RBS were simulated and compared to the existing simulation results (from **Figures S8** and **S10**) performed for equivalent long CDS constructs. Un‐ optimised describes a CDS with some poor codons interspersed throughout the sequence. De-optimised describes intentional introduction of many poor codons at one point in the sequence in order to create a potential translational bottleneck. **(B)** The capacity and estimated construct output produced by short CDS (846 bp) scFv constructs F1, F2, and F5 with 1 hour post‐induction in DH10B cells in exponential growth. Existing data from Figure 2A for equivalent long CDS VioB-mCherry constructs H2, H5 and H4 are also shown for comparison. Bold text indicates the use of strong RBS sequences. Filled bars indicate induced samples and hatched bars uninduced. Capacity and growth rate are measured as for Figure 2A. scFv output is estimated from the band intensity of the Western blot shown in Figure S16. Error bars represent the standard deviation of 3 independent repeats.

**Figure S15:**
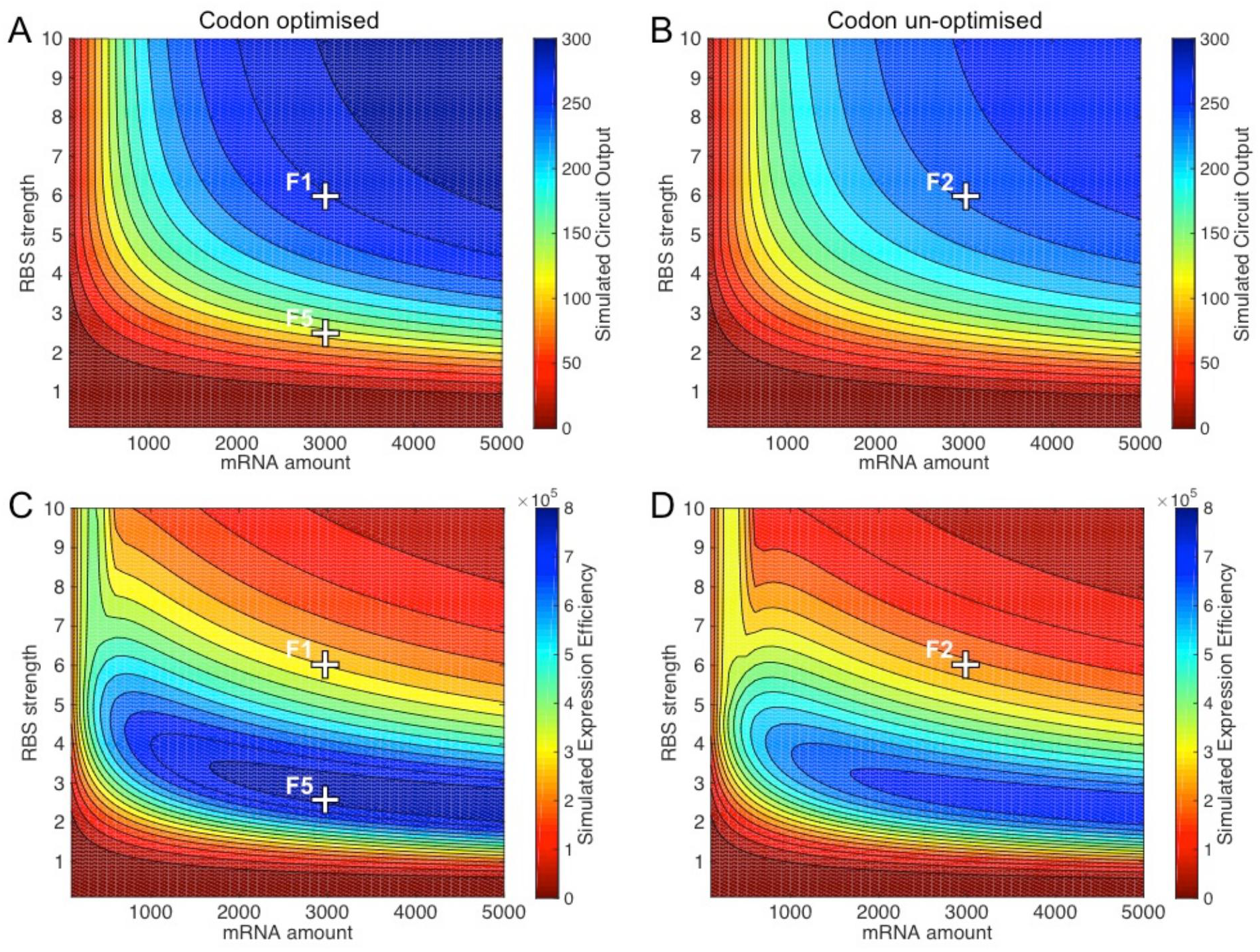
Heatmap of simulated construct outputs for codon optimised (**A**) and un‐optimised **(B**) short CDS constructs alongside heatmaps of simulated expression efficiency for codon optimised (**C**) and un‐optimised (**D**) constructs. Locations of simulations mapping to all constructs assayed in Figure S14 are shown. Simulated circuit output is the simulated production rate of the circuit protein at steady state. Simulated expression efficiency is calculated as the product of circuit output and number of free ribosomes available at steady state.

**Figure S16:**
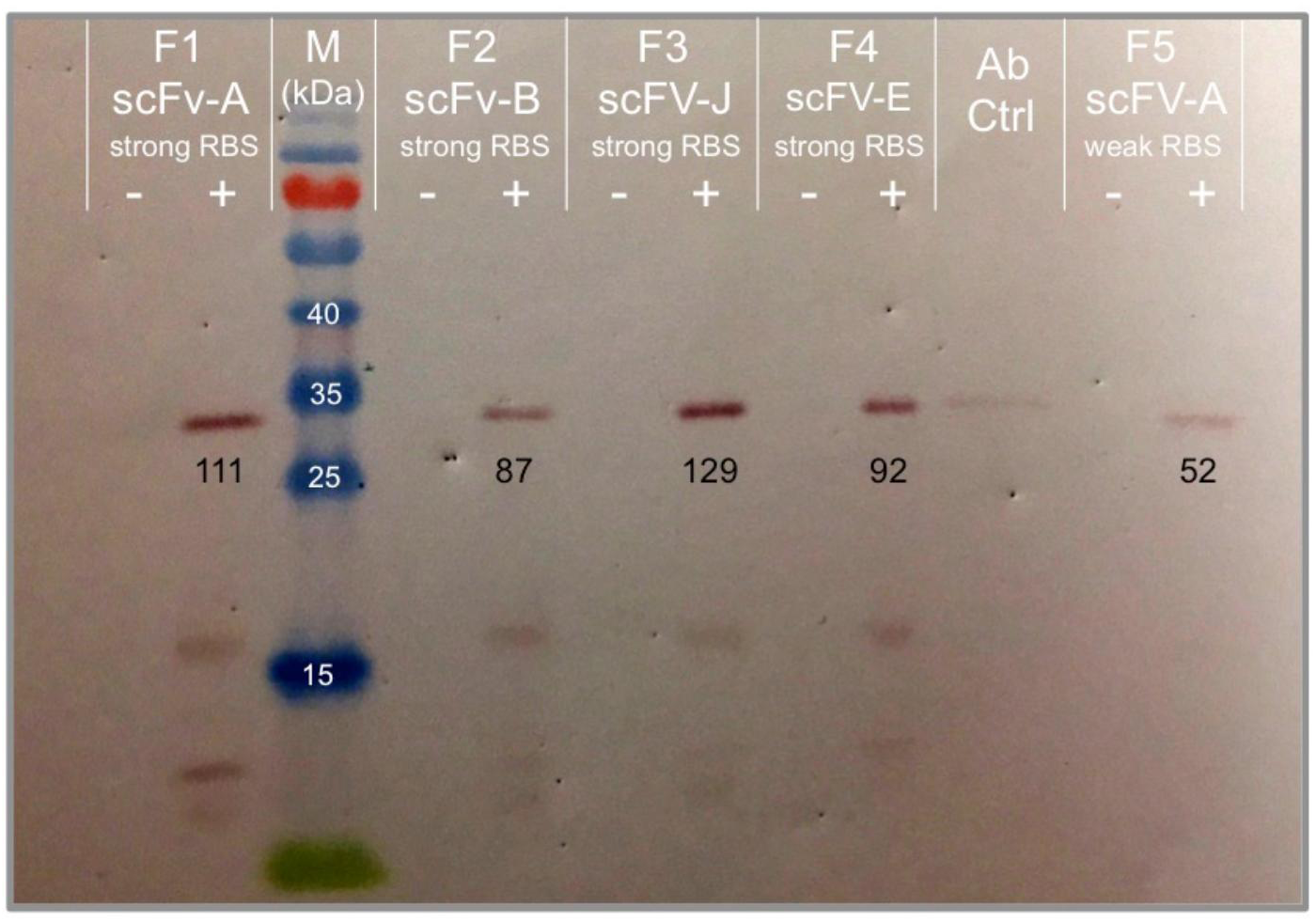
Western blot of DH10B*E. coli*with capacity monitor expressing constructs F1, F2, F3, F4 and F5 with (+)and without (‐) construct induction. Protein was harvested from cell culture 4 hours post‐induction. Blot shows expression of His-tagged scFv constructs (32 kDa) as well as the antibody positive control (Ab Ctrl) and 10-170 kDa size marker (M) separated by SDS‐PAGE (12%). scFC expression was estimated by subtracting band intensity compared from surrounding background intensity using ImageJ. Intensity measurements are given on the gel in black text below each band. Size marker molecular weights (kDa) are given in white text.

**Figure S17:**
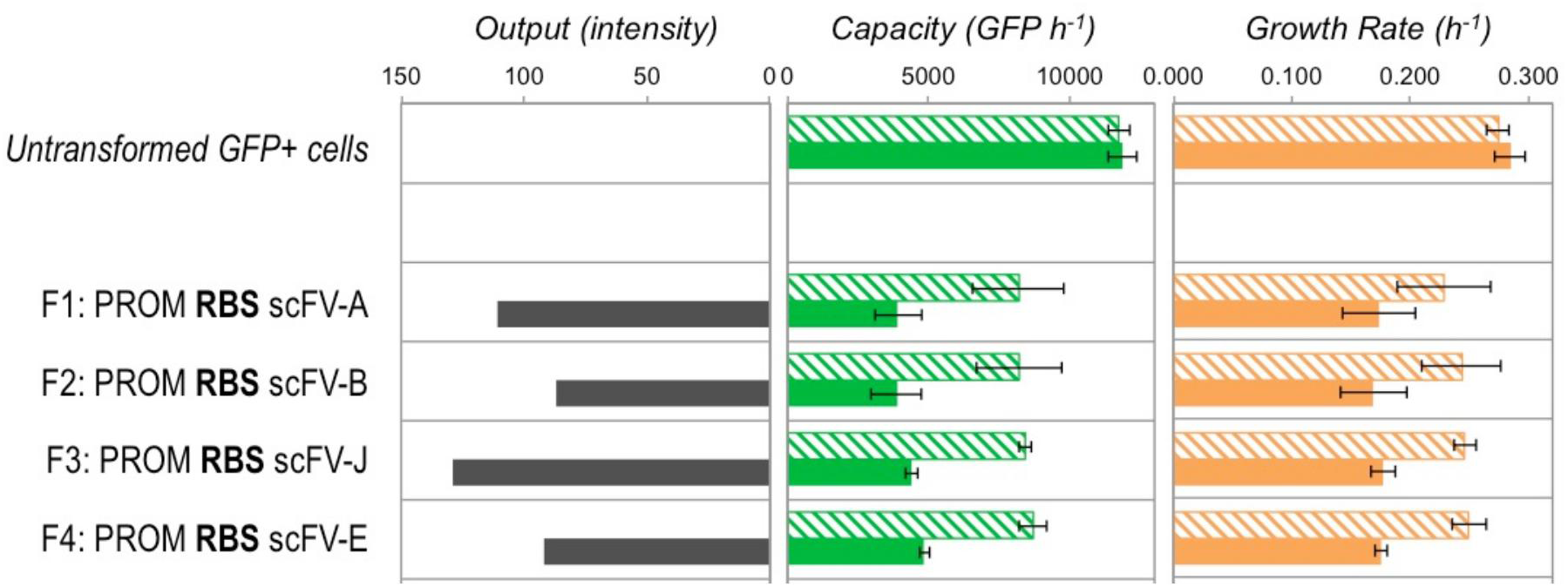
The capacity, growth rate and estimated construct output produced by scFv constructs F1, F2, F3 and F4with different codon optimisation 1 hour post-induction in DH10B cells in exponential growth. This figure is discussed in **Supplementary Note 4**. Bold text indicates the use of strong RBS sequences. Filled bars indicate induced samples and hatched bars uninduced. Capacity and growth rate are measured as for Figure 2A, output is estimated from the band intensity of the Western blot shown in Figure S16. Error bars represent the standard deviation of 3 independent repeats.

**Table S1.:**
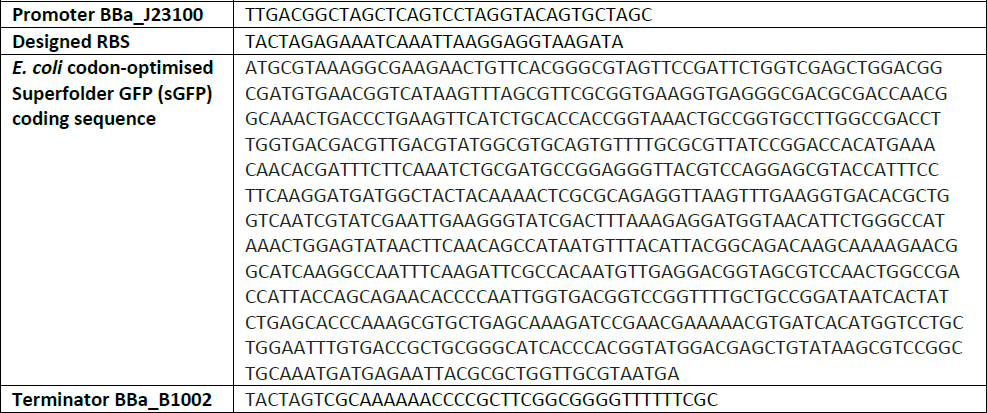
Sequence of the synthetic parts encoding the capacity monitor cassette. Sequences for the BBa_J23100 constitutive promoter and BBa_B1002 terminator were obtained from the iGEM Parts Registry while the RBS sequence was designed with the RBS calculator^32^. The superfolder GFP coding sequence^44^ was designed by DNA2.0 (Menlo Park, CA) for efficient high-level expression in *E. coli.*

**Table S2.** Diagrams of the Luc, Lux and sCG inducible constructs used for the current study. The firefly luciferase Luc operon, the LuxCDABEG operon from *Vibrio fischeri* and the human Chorionic Gonadrophin beta subunit (sCG) are all codon-optimised for *E. coli* and regulated by the L-arabinose-inducible AraC-pBad regulator-promoter cassette and are on pSB1C3 high-copy plasmid with chloramphenicol selection.

| Construct Name | Schematic Representation |
| --- | --- |
| AraC-pBAD + Optimised Firefly Luciferase Operon [Luc] on pSB1C3 |  |
| AraC-pBAD + Optimised *V. Fischeri* Luciferase Operon [Lux] on pSB1C3 |  |
| AraC-pBAD + *E. coli* optimised Chorionic Gonadotrophin [sCG] on pSB1C3 |  |

**Table S3.**
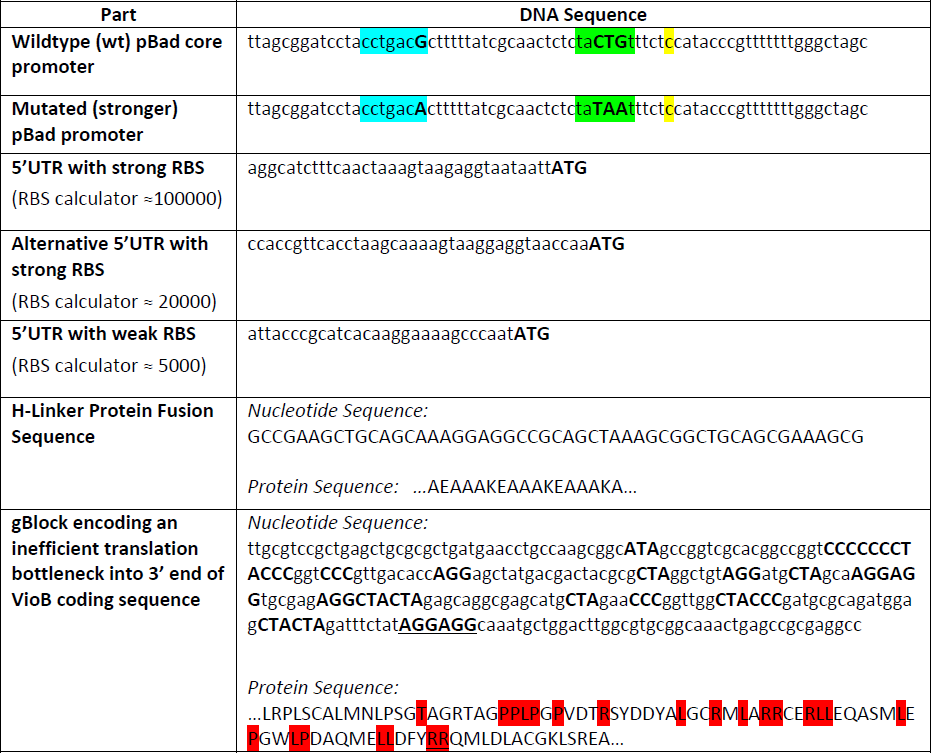
Sequences of designed parts used for the construction of the VioB‐mCherry combinatorial library. The modified pBad core promoter sequence was described by the 2011 DTU Denmark iGEM team to increase expression from pBad. Point mutations are shown in bold uppercase and the -35 and ‐10 and +1 sequences are highlighted in green, blue and yellow respectively. VioB 5’UTR regions were designed to incorporate RBS sequences generated by the RBS calculator (lowercase) in forward engineering mode^32^. The H‐linker sequence was previously shown to be a suitable linker for bifunctional fusion protein^31^. The 267 bp gBlock was designed to replace optimised *E. coli* codons at the 3’end of the VioB coding sequence with inefficient coding by introducing synonymous mutations (bold uppercase) that incorporate rare codons and a strong anti‐Shine Dalgarno sequence^38^. These are shown in red and underlined respectively in the protein sequence.

**Table S4:** Part composition of transcription factor‐regulated constructs used in Figures S3 & S4, with BioBrick numbers denoting the parts as listed in the iGEM Parts Registry. The LacI system construct was assembled from parts for this study. The LuxR system was already available as a complete construct (BBa_F2620). The TetR system is an abridged version the TetR-mRFP, already available as the complete BBa_I732913 construct.

| Regulator | Part Composition (including BioBrick numbers) |  |  |  |  |  |  |  |  |
| --- | --- | --- | --- | --- | --- | --- | --- | --- | --- |
| **LacI system**<br>isopropyl β-D-1-thiogalactopyranoside (IPTG) | plasmid<br>pSB1A2 | promoter<br>J23105 | RBS<br>B0034 | Regulator CDS<br>LacI<br>C0012 | terminator<br>B0015 | promoter<br>pLac<br>R0010 | | | |
| **LuxR system**<br>acylhomoserinelactone (AHL)<br>**BBa_F2620** | plasmid<br>pSB1C3 | promoter<br>p(tetR)<br>R0040 | RBS<br>B0034 | Regulator CDS<br>LuxR<br>C0062 | terminator<br>B0015 | promoter<br>pLux<br>R0062 |  |  |  |
| **TetR system**<br>anhydrotetracycline (aTc) | plasmid<br>pSB1A2 | promoter<br>p(lacIQ)<br>I14032 | RBS<br>B0034 | Regulator CDS<br>TetR<br>I732080 | terminator<br>B0015 | promoter<br>p(tetR)<br>R0040 |  |  |  |
| **TetR-mRFP system**<br>anhydrotetracycline (aTc)<br>**BBa_I732913** | plasmid<br>pSB1A2 | promoter<br>p(lacIQ)<br>I14032 | RBS<br>B0034 | Regulator CDS<br>TetR<br>I732080 | terminator<br>B0015 | promoter<br>p(tetR)<br>R0040 | RBS<br>B0034 | Reporter CDS<br>mRFP1<br>E1010 | terminator<br>B0015 |

**Table S5.**
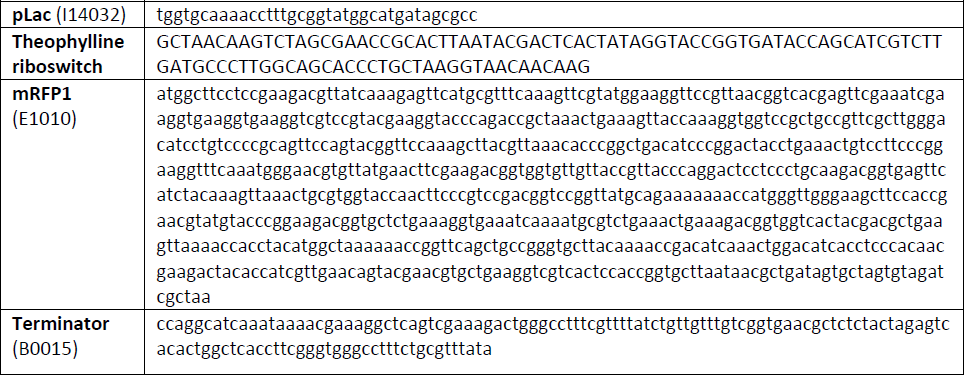
Components of the theophylline-induced riboswitch system used in Figure S4, with BioBrick numbers denoting the parts as listed in the iGEM Parts Registry. The theophylline riboswitch sequence was obtained from a previous study^29^.

**Table S6.**
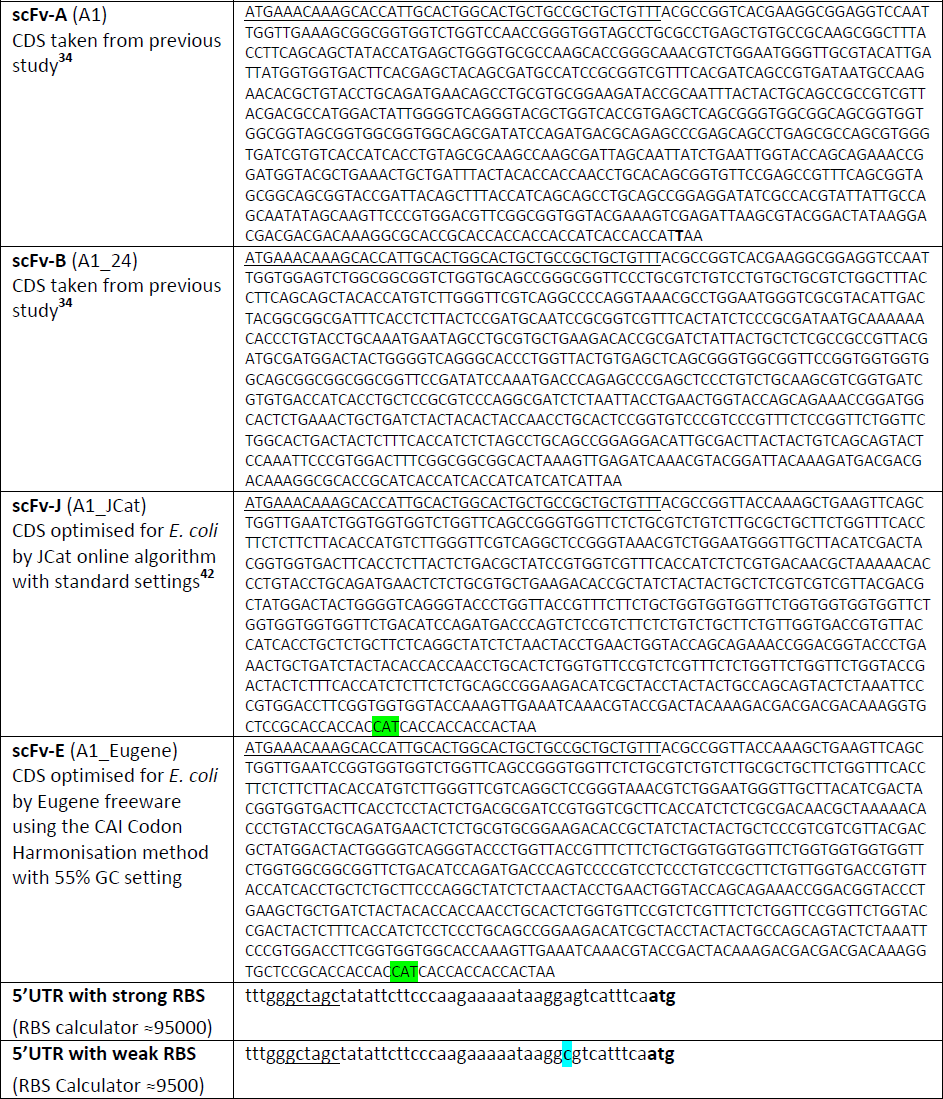
Sequences of parts in scFV constructs F1 to F5 used in Figures S14, S16 & S17. All constructs contained AraC‐ pBad as the upstream regulator/promoter and were on high‐copy pSB1C3 plasmid. For all scFv CDS sequences, the first 45 bp are underlined and are kept the same. For both scFv‐J and scFv-E sequences, a CAT codon (green) was manually swapped in place of a CAC codon in one position to remove prohibitive DNA sequence repetition at the C-terminal HisTag. The weak RBS was generated by a point mutation (blue) in the strong RBS. An NheI digest site (underlined) in both RBS sequences facilitated construct cloning

## TRANSLATION MODEL

A model of how a synthetic gene and native host cell processes compete for shared resources was constructed by focussing on the process of translation, specifically the use of ribosomes. In order to create a model that could capture some of the key processes, we looked carefully at the process of elongation, as ribosomes spend a majority of the translational process in this stage. In addition, it is well known that different profiles of elongation rates can cause phenomena such as ‘traffic jams’ that can only be explained when individual elongation rates are taken into consideration^36, 38^.

The assumptions and derivation details for the proposed translation model can be found in our companion theoretical paper^18^.

## DEVELOPING A SIMULATION FOR RIBOSOME USE DURING TRANSLATION

A Python script was built that allowed our model to be simulated. It consists of two classes, Circuit and Cell. The Circuit class describes gene expression constructs and allows a user to define the number of transcripts, elongation rates, and the binding and unbinding affinities for the RBS. This can be done for any number of constructs. The Cell class allows a user to define a model of a cell including the total number of ribosomes available in the cell as well as which construct(s) it contains. These classes have attributes and methods that allow simulations of the system to be run. The method simulated on the Cell class allows the simulation of the cell to be run and gives a dictionary output that describes the number of free ribosomes remaining in the cell as well as the distribution of ribosomes for each circuit. In a modification to the published model^18^, elongation here is defined as a series of steps of 10 codons (30 nucleotides) at a time, rather than one codon at a time, as this is the footprint of each ribosome on an mRNA and better represents how many can be queued on a transcript. This script uses functions provided by the scipy Python package, which thus must be installed *a priori*

The content of the Python file implementing the translation model described above (called models.py in our Python package) is copied hereafter for the interested reader.

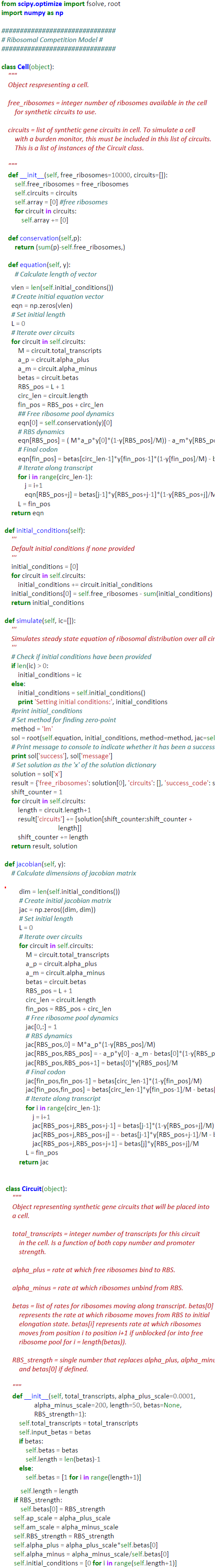

***Model simulations with realistic parameter values***. In order to test our model, we performed an initial numerical simulation of output from a genomic gene with realistic values for *E. coli* obtained from Bionumbers.org. The table below shows the parameters used during this simulation. These parameters roughly correspond to one copy of a gene with a medium-strength promoter (3 transcripts per promoter in a cell at any time) and optimised coding sequence (elongation rate = 1 for all steps). The coding sequence is set as 50 steps, which is equivalent to a 1500 bp (500 amino acids, or 50 codon groups) gene with steps equal to 30 nucleotides.

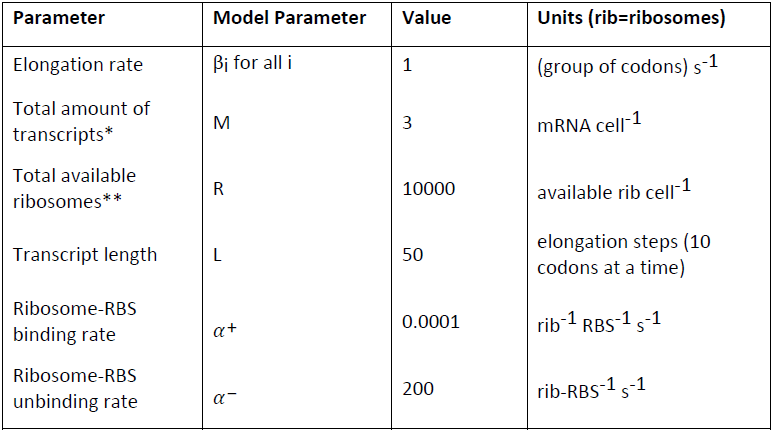
Model Parameters Used For Testing Model Validity

* For a medium strength promoter (estimated from Bionumber ID 107667)
Estimate for ribosomes per cell at 37° C with doubling time of 40 mins is 26300 (Bionumber ID 102015), but in exponentially growing *E. coli* the max percentage of unnecessary protein expression is 30 to 40% ^14^ and therefore 38% of total ribosomes (10000) are available for this simulation.

Running this simulation shows that the gene uses an average of 41.8 ribosomes at any point in time, which is 0.16% of all cellular ribosomes, assuming the total number of ribosomes is 26300 (Bionumber ID 102015). This appears to be the correct order of magnitude since there are approximately 4000 genes in the cell, of which around half are active in exponential growth. This gives 2000 active promoters with, on average, 1‐5 mRNA transcripts per promoter per cell, giving a total of 2000‐10000 mRNA transcripts per cell. The 3 mRNA transcripts from the simulated gene constitute 0.03‐0.15% of the total cellular mRNAs, and therefore it appears valid that these transcripts use 0.16% of the cell’s ribosomes as they have a medium‐strength RBS.

## ACCOMPANYING PYTHON PACKAGE FOR SIMULATING A LANDSCAPE OF CONSTRUCT DESIGNS

To make the simulation of a landscape of construct designs differing by the various tuning knobs accessible experimentally, e.g. the amount of mRNA (adjustable by promoter strength and/or plasmid copy‐number), the RBS strength, and the translation rate (adjustable by codon choice), we created a self‐standing Python package.

This package should be unzipped and, using the command line, the user should navigate into the unzipped directory. Once in the directory, the user can run the simulation using the following command: python simulate_landscape.py [xml_location] where [xml_location] is to be replaced by the name of the parameter‐defining xml file (if in the same directory, e.g. fast_codons_parameters.xml), or the location and name of the xml file (e.g. /Users/username/synbio_simulations/parameter_files/fast_codon_paramters.xml). This will then output the data into separate csv files. Each row in these csv files contains data corresponding to a single RBS strength and each column contains data corresponding to a single number of mRNA. The top left cell contains data associated with the lowest RBS strength and lowest number of mRNA with the number of RBS increasing by rbs_strength_step in each subsequent row and the number of mRNA increasing by mrna_number_step in each subsequent column.

The unzipped package contains the following files:

- simulate_landscape.py
- simulations.py
- models.py

and a directory called data where the simulation data are deposited.

The contents and set of commands needed to use these Python files is explained in the accompanying README.txt file. As a result of the execution of the file simulate_landscape.py various .csv files are created within the directory data:

- circuit_out.csv
  - This file contains the protein production rates per cell for the synthetic circuit.
- circuit_ribs.csv
  - This file contains the total numbers of ribosomes sequestered on all of the transcripts for the synthetic circuit.
- free_ribs.csv
  - This file contains the numbers of free ribosomes per cell.
- monitor_out.csv
  - This file contains the protein production rates per cell for the burden monitor.
- monitor_ribs.csv
  - This file contains the total numbers of ribosomes sequestered on all of the transcripts for the burden monitor.
- mRNA_amount.csv
  - This file contains a vector of all the mRNA amounts used in the simulations across the landscape.
- RBS_str.csv
  - This file contains a vector of all the RBS strengths used in the simulations across the landscape.

## VISUALISATION OF THE LANDSCAPE SIMULATION RESULTS

The generated data contained in the .csv files listed above can be visualised using various programs, e.g. Microsoft Excel, GraphPad Prism, Matlab, etc. To make the visualisation of “efficiency” easier, we also created a Matlab script called Plot_Matlab_3D_CSV.m. This script needs to be placed in the same directory as the one containing the various .csv files listed above. Once called in Matlab, the execution of this script will create a PDF file calledHeat_Map_Efficiency.pdf representing the Normalised Efficiency, which is calculated as the product of circuit output and number of free ribosomes, normalised to the maximum efficiency value (see Figure 2D, **S10** & **S15**).

## Acknowledgements

The authors wish to thank James J. Collins, Matthew Scott, Dan Goodman, Anil Wipat, Dan Siegal‐Gaskins, Julius Lucks, James Chappell, Chris Hirst, Kate Royle and CSYNBI colleagues for thoughts and advice during this project. This work was supported by grants from the UK Engineering and Physical Research Council (EP/G036004/1, EP/J021849/1 and EP/J02175X/1).

